# The transcription and export complex THO/TREX contributes to transcription termination in plants

**DOI:** 10.1101/2020.03.02.972356

**Authors:** Ghazanfar Abbas Khan, Jules Deforges, Rodrigo S. Reis, Yi-Fang Hsieh, Jonatan Montpetit, Wojciech Antosz, Luca Santuari, Christian S Hardtke, Klaus Grasser, Yves Poirier

**Affiliations:** Department of Plant Molecular Biology, University of Lausanne, Switzerland; School of Biosciences, University of Melbourne, VIC, Australia; Department of Cell Biology & Plant Biochemistry, Biochemistry Centre, University of Regensburg, Universitätsstr. 31, D-93053 Regensburg, Germany

## Abstract

Transcription termination has important regulatory functions, impacting mRNA stability, localization and translation potential. Failure to appropriately terminate transcription can also lead to read-through transcription and the synthesis of antisense RNAs which can have profound impact on gene expression. The Transcription-Export (THO/TREX) protein complex plays an important role in coupling transcription with splicing and export of mRNA. However, little is known about the role of the THO/TREX complex in the control of transcription termination. In this work, we show that two proteins of the THO/TREX complex, namely TREX COMPONENT 1 (TEX1 or THO3) and HYPER RECOMBINATION1 (HPR1 or THO1) contribute to the correct transcription termination at several loci in *Arabidopsis thaliana*. We first demonstrate this by showing defective termination in *tex1* and *hpr1* mutants at the nopaline synthase (NOS) terminator present in a T-DNA inserted between exon 1 and 3 of the *PHO1* locus in the *pho1-7* mutant. Read-through transcription beyond the NOS terminator and splicing-out of the T-DNA resulted in the generation of a near full-length *PHO1* mRNA (minus exon 2) in the *tex1 pho1-7* and *hpr1 pho1-7* double mutants, with enhanced production of a truncated PHO1 protein that retained phosphate export activity. Consequently, the strong reduction of shoot growth associated with the severe phosphate deficiency of the *pho1-7* mutant was alleviated in the *tex1 pho1-7* and *hpr1 pho1-7* double mutants. Additionally, we show that RNA termination defects in *tex1* and *hpr1* mutants leads to 3’UTR extensions in several plant genes. These results demonstrate that THO/TREX complex contributes to the regulation of transcription termination.

**Author summary:** Production of messenger RNAs (mRNAs) involves numerous steps including initiation of transcription, elongation, splicing, termination, as well as export out of the nucleus. All these steps are highly coordinated and failure in any steps has a profound impact on the level and identity of mRNAs produced. The THO/TREX protein complex is associated with nascent RNAs and contributes to several mRNA biogenesis steps, including splicing and export. However, the contribution of the THO/TREX complex to mRNA termination was poorly defined. We have identified a role for two components of the THO/TREX complex, namely the proteins TEX1 and HPR1, in the control of transcription termination in the plant *Arabidopsis thaliana*. We show that the *tex1* and *hpr1* mutants have defects in terminating mRNA at the nopaline synthase (NOS) terminator found in T-DNA insertion mutants leading to the transcriptional read-through pass the NOS terminator. We also show that *tex1* and *hpr1* mutants have defects in mRNA termination at several endogenous genes, leading to the production of 3’UTR extensions. Together, these results highlight a role for the THO/TREX complex in mRNA termination.

## Introduction

In eukaryotes, mRNAs are generated by several dynamic and coordinated processes including transcriptional initiation, elongation and termination, as well as splicing and nuclear export. Failure in any of these processes has a profound impact on the level and identity of transcripts [1–3]. These transcriptional steps are sequentially orchestrated by a multitude of RNA-binding protein complexes that co-transcriptionally couple with the nascent RNA [4]. For example, protein complex required for transcription termination cleaves pre-mRNA close to RNA polymerase II (RNAPII) and adds a poly (A) tail to the 3’end of nascent RNA [3]. Pre-mRNA cleavage exposes 5’end of nascent mRNA to 5’-3’ exonucleases which degrades the RNA attached to RNAPII, leading to transcription termination [5].

Cleavage and polyadenylation define the transcription termination at a given locus and has a decisive role in regulating gene expression as it can influence stability and translation potential of the RNA via the inclusion of regulatory sequence elements [6]. Moreover, transcription termination avoid interference with the transcription of downstream genes and facilitates RNAPII recycling [7]. It also prevents synthesis of antisense RNAs which can have a severe effect on RNA production and overall gene expression [8]. Additional regulatory role of transcription termination is the synthesis of chimeric transcripts formed by tethering of two neighboring genes on the same chromosomal strand [8]. Considering its importance, molecular mechanisms which regulate RNA termination are relatively poorly understood.

After transcription termination, nascent RNA is assembled in a ribonucleoprotein complex is delivered for RNA export into the cytosol. Similar to other steps in RNA biogenesis which are closely coupled in a sequential manner, it is likely that transcription termination is associated with nuclear export of RNAs. The TREX (TRanscription-EXport) protein complex has emerged as an important component in coupling transcription with RNA processing and export [9]. In metazoans, TREX consists of THO core complex, which includes THO1/HPR1, THO2, THO3/TEX1, THO5, THO6 and THO7 [10]. The proteins associating with the THO core components and forming the TREX complex include the RNA helicase and splicing factor DDX39B (SUB2 in yeast) as well as the RNA export adaptor protein ALY (YRA1 in yeast) [10]. TREX is co-transcriptionally recruited to the nascent mRNA and regulates splicing, elongation and export [11]. Moreover, THO/TREX complex is required for the genetic stability as it is required for preventing DNA:RNA hybrids that lead to transcription impairment and are responsible for genetic instability phenotypes observed in these mutants [12]. The genes encoding the THO core components are conserved in plants, including in the model plant *Arabidopsis thaliana* [13, 14]. *A. thaliana* mutants defective in THO components show a wide range of phenotypes, from no obvious alteration and relatively mild phenotypes to lethal phenotypes, suggesting overlapping but independent functions of these components [14–16]. *A. thaliana* mutants in THO components, including HPR1, THO2 and TEX1, show defects in small RNA biogenesis, mRNA elongation, splicing and export, but no defect in mRNA termination has been reported [14–20].

In this work, we explored the regulatory function of two components of the THO/TREX complex, namely TEX1 and HPR1, in transcription termination in *A. thaliana*. We show that the *tex1* and *hpr1* mutants are defective in RNA termination at the Nopaline Synthase (NOS) terminator present on a T-DNA inserted in the *PHO1* locus. Additionally, genome-wide analysis of mRNAs revealed RNAPII termination defects in *tex1* and *hpr1* mutants at several loci leading to the 3’UTR extensions.

## Results

### A forward genetic screen using the T-DNA insertion mutant *pho1-7* for reversion of the growth phenotype identified the *TEX1* gene

The *PHO1* gene encodes an inorganic phosphate (Pi) exporter involved in loading Pi into the root xylem for its transfer to the shoot [21–24]. Consequently, *pho1* mutants in both *A. thaliana* and rice have reduced shoot Pi contents and shows all symptoms associated with Pi deficiency, including highly reduced shoot growth and the expression of numerous genes associated with Pi deficiency [23, 25, 26]. However, it has previously been shown that low shoot Pi content can be dissociated from its major effects on growth and other responses normally associated with Pi deficiency through the modulation of *PHO1* expression or activity [25, 27]. We thus used the *pho1* mutant as a tool, in a forward genetic screen, to identify mutants which restore *pho1* shoot growth to wild type (Col-0) level, while maintaining low shoot Pi contents. We performed ethyl methane sulphonate (EMS) mutagenesis on seeds of the *pho1-7* mutant, derived from the SALK line 119520 containing a single T-DNA inserted in the *PHO1* gene in between the first and third exon of the *PHO1* gene leading to the deletion of the second exon from the genome (S1 Text). Screening of 10’300 mutagenized *pho1-7* plants grown in soil for improved rosette growth lead to the isolation of a suppressor mutant and had rosette growth similar to Col-0 while maintaining a low shoot Pi content similar to *pho1-7* (Fig 1AB) (see Material and methods for further detail). Both *pho1-7* and the suppressor mutant maintained resistance to kanamycin associated with the T-DNA insertion in *PHO1*.

Mapping-by-sequencing revealed that the mutation C116T is introduced into the *TEX1* gene in the *pho1-7* suppressor mutant. This leads to a conversion of amino acid serine 39 to phenylalanine in the TEX1 protein. Transformation of *TEX1* gene into *pho1-7* suppressor led to *pho1*-like phenotype, confirming that mutation in *tex1* was the causal mutation for restoration of *pho1-7* shoot growth (Fig 1AB). Furthermore, crossing of a T-DNA allele *tex1-4* (SALK_100012) to *pho1-7* also resulted in the suppression of *pho1-7* shoot growth phenotype (Fig 2A), further confirming that mutation in *TEX1* is responsible for the suppression of shoot growth phenotype in *pho1-7 suppressor* mutant. Therefore, we named this new S39P mutant in the *TEX1* gene as *tex1-6*. In agreement with previous reports, TEX1 protein was localized to the nucleus [19] (S1A Fig). *TEX1* promoter fusion with GUS showed that *TEX1* is expressed in root, cotyledon and rosette (S1B Fig).

The *pho1-7 tex1-6* mutant shows Col-0-like shoot growth while maintaining a low Pi content that is only slightly higher to the parental *pho1-7* (Fig 1A, B). A key molecular response of *pho1* mutants is the manifestation of gene expression and lipid profiles in the shoots that are associated with strong Pi deficiency [25]. To determine if the *tex1-6* mutation can also suppress the induction of Pi starvation responses (PSR) in the rosettes of *pho1-7 tex1-6*, we performed quantitative RT-PCR (qRT-PCR) to see the expression of PSR genes. *pho1-7 tex1-6* shoots showed an expression profile of PSR genes that was comparable to Pi-sufficient Col-0 plants (S2A Fig). Additionally, lipid analysis in *pho1-7* mutants showed a decrease of phospholipids and an increase in galactolipids expected for Pi-deficient plants, while *pho1-7 tex1-6* plants showed lipids profiles similar to Col-0 plants (S2B Fig), confirming that the *tex1-6* mutation suppressed morphological as well as molecular response to Pi deficiency displayed by the *pho1-7* mutant.

**Fig 1.**
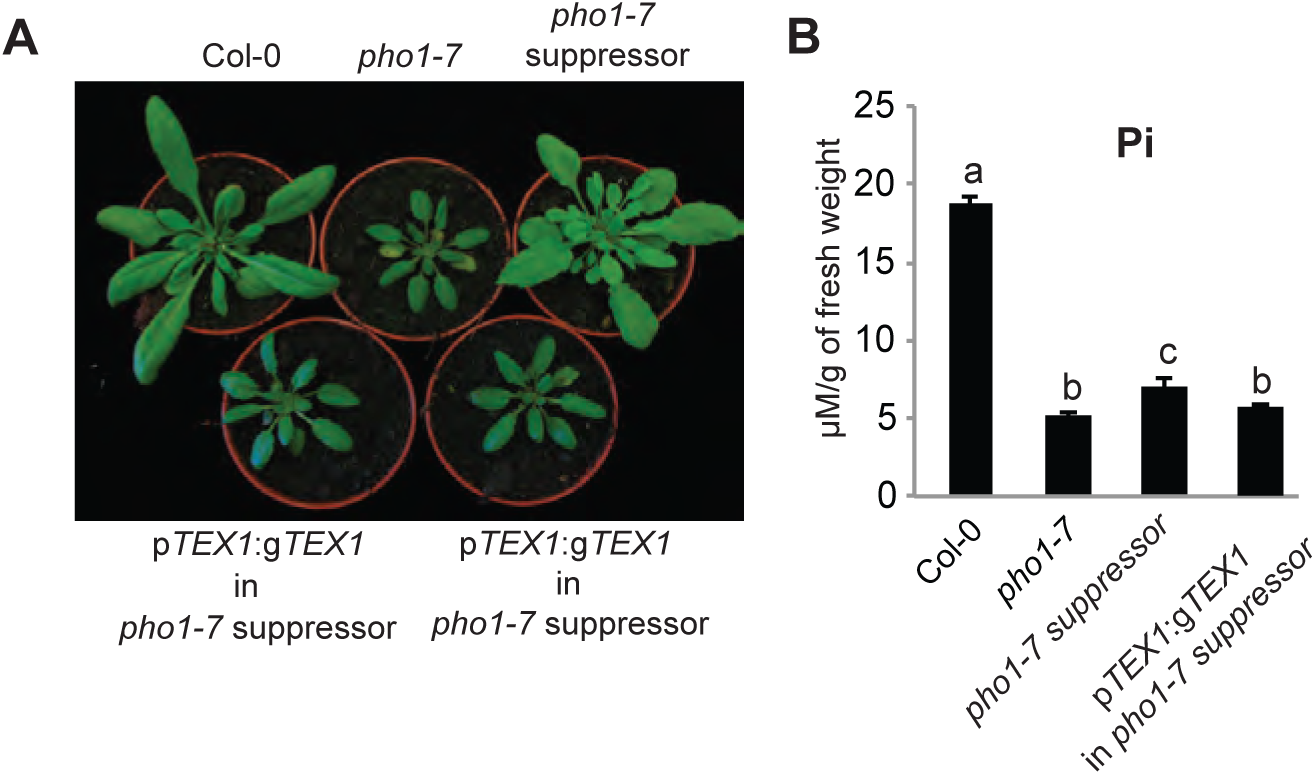
The *tex1* mutation suppresses the shoot growth phenotype of the *pho1-7* mutant. (A) Phenotype of Col-0, *pho1-7*, *pho1-7 suppressor*, and *pho1-7 suppressor* transformed with the p*TEX1:gTEX1:GFP*. (B) Pi contents of 4-week-old rosettes. Data from a representative experiment shows the mean Pi content from ten different plants grown in independent pots. Errors bars represent standard deviation. Values marked with lowercase letters are statistically significantly different from those for other groups marked with different letters (P < 0.05, ANOVA with the Tukey-Kramer HSD test). Panel display a representative experiment

### Mutation of *TEX1* in *pho1-7* resulted in the synthesis of a truncated PHO1 protein

To determine if the *tex1* mutation also suppresses the growth phenotype associated with other *pho1* alleles generated by EMS mutagenesis, a double mutant *pho1-4 tex1-4* was generated. Surprisingly, *pho1-4 tex1-4* double mutant showed only minor improvement in shoot growth and maintained low shoot Pi content (Fig 2A, B). In order to understand how the *tex1* mutation can result in restoration of *pho1-7* shoot growth, we performed a detailed analysis of transcripts produced at the *PHO1* locus in the *pho1-7 tex1-4* mutant. Interestingly, we identified a truncated *PHO1*^Δ249-342^ transcript which only lacked the 2^nd^ exon suggesting that the T-DNA is spliced out from the mature mRNA (Fig 2C, S1 Text). The mRNA produced is in frame and resulted in the production a truncated PHO1^Δ84-114^ protein (S1 Text). Western blot experiments confirmed the presence of a PHO1^Δ84-114^ truncated protein in both *pho1-7* and *pho1-7 tex1-4* roots with a strong increase of expression in the *pho1-7 tex1-4* double mutant as compared to *pho1-7* (Fig 2D). This increase in protein quantity can be attributed to an increase in expression of *PHO1*^Δ249-342^ RNA in the *pho1-7 tex1-4* double mutant (Fig 2E). We hypothesized that the PHO1^Δ84-114^ protein variant was at least partially active as a Pi exporter and that its increased expression in *pho1-7 tex1-4* partially restored PHO1 function, resulting in an improvement of the shoot growth phenotype. We confirmed this hypothesis by expressing the *PHO1^Δ84-114^* variant and the wild type *PHO1* fused to GFP using the *PHO1* promoter in the *pho1-4* null mutant. As expected, *pho1-4* mutant which expressed the wild type PHO1 fully complemented the growth and Pi content to Col-0 level (Fig 3A, B). However, plants expressing the PHO1^Δ84-114^ variant only restored the shoot growth phenotype while maintaining low Pi contents comparable to *pho1-7 tex1-4* plants (Fig 3A, B). Confocal analysis of roots showed that PHO1^Δ84-114^ variant protein was localized similarly to wild type PHO1 (Fig 3C), which was previously shown to be primarily in the Golgi and *trans*-Golgi network (TGN) [21]. Furthermore, transient expression of the PHO1^Δ83-114^-GFP fusion in *Nicotiana benthamiana* leaves led to Pi export to the apoplastic space, demonstrating that the protein was competent in Pi export (Fig 3D) [21]. Collectively, these results confirmed that restored expression of *PHO1*^Δ249-342^ RNA is responsible for the improved shoot growth in *pho1-7 tex1-4* double mutants.

**Fig 2.**
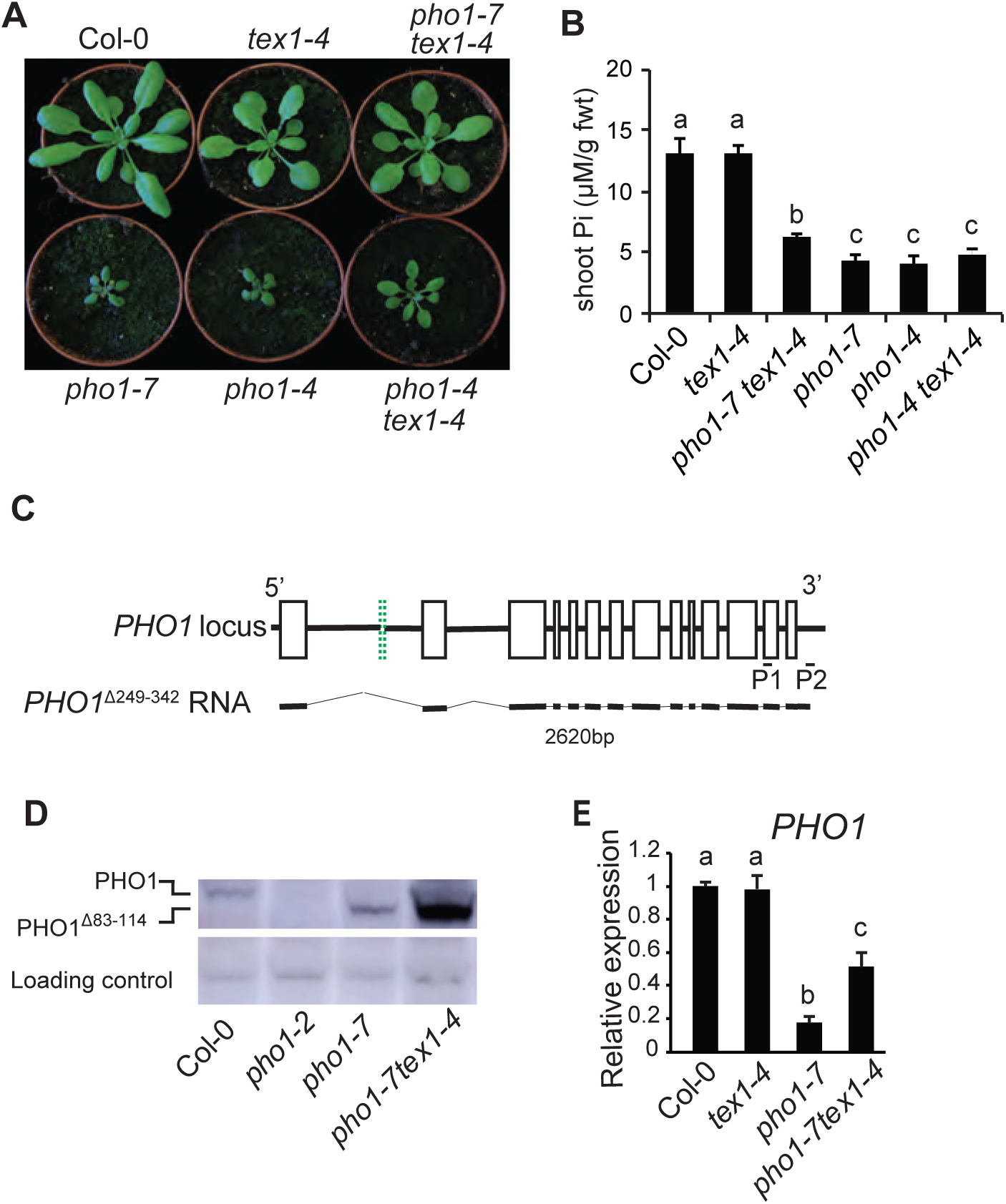
*tex1* restores the expression of PHO1 in the *pho1-7* mutant. (A) Phenotype of 4-week-old Col-0, *tex1-4*, *pho1-7*, *pho1-4*, *pho1-7 tex1-4* and *pho1-4 tex1-4* mutants. (B) Shoot Pi contents of 4-week-old plants. Data from a representative experiment shows the means of Pi contents from six individual plants grown in independent pots. Error bars represent standard deviation. (C) Structure of truncated *PHO1* mRNA produced at the *PHO1* locus in the *pho1-7* mutant. For the *PHO1* locus, exons are indicated as black open boxes, except for the second exon, indicated as a green doted box. Exon 2 is present in Col-0 but deleted in the *pho1-7* mutant as a result of T-DNA insertion. The structure of the mRNA *PHO1^Δ249-342^* found in the *pho1-7* mutant is shown below. The pairs of oligonucleotides P1 and P2 used for RT-PCR is shown. (D) Western blot showing full length and truncated PHO1 protein in roots of Col-0, *pho1-2* null mutant, *pho1-7* and *pho1-7 tex1-4*. Plants were grown for 4 weeks in a clay substrate and total protein were extracted from roots. (E) Relative expression level of *PHO1* gene in the roots of 4-week-old plants grown in a clay substrate. Data are means of three samples from plants grown in independent pots and three technical replicates for each sample, with each sample being a pool of three plants. Error bars represent standard deviation. For both B and E, values marked with lowercase letters are statistically significantly different from those for other groups marked with different letters. (P < 0.05, ANOVA with the Tukey-Kramer HSD test).

**Fig 3.**
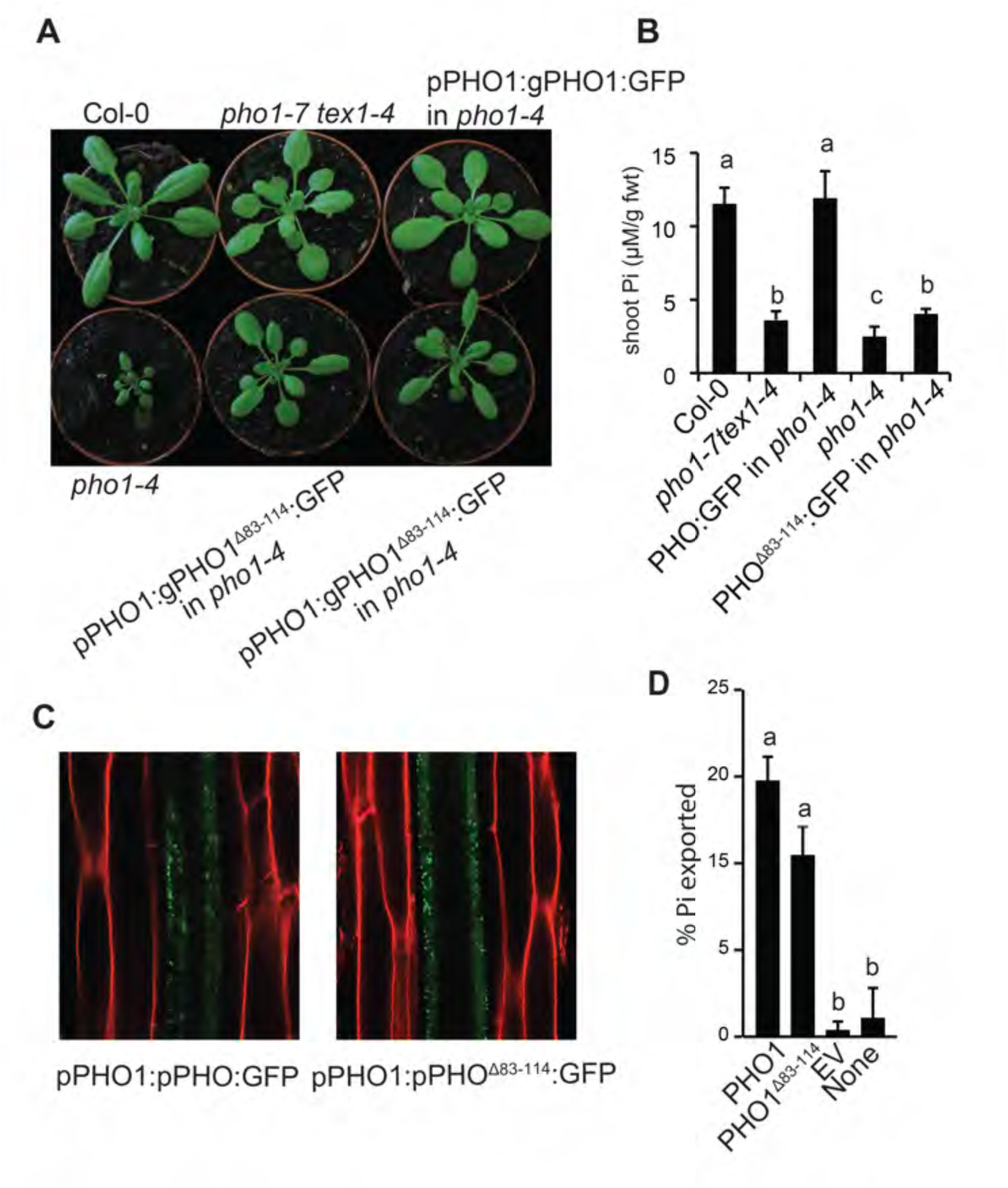
Truncated PHO1^Δ84-114^ is sufficient to restore the shoot growth phenotype of *pho1-4* null mutant. (A) Phenotype of four-week-old plants. (B) Pi contents in the rosettes of 4-week-old plants. Data from a representative experiment shows the means of Pi contents from six individual plants grown in independent pots. Error bars represent standard deviation. (C) Localization of full-length PHO1:GFP and PHO1^Δ83-114^:GFP in the roots of 7-day-old seedlings. Both *PHO1:GFP* and *PHO1^Δ83-114^:GFP* were expressed under the control of native *PHO1* promoter in the *pho1-4* null mutant. (D) Pi export mediated by PHO1:GFP and PHO1^Δ84-114^:GFP from transient expression in *N. benthamiana* leaf discs. As controls, Pi export was measured in leaf discs expressing either free GFP (EV) or not infiltrated (none). Data are means of 4 measurements taken from independent infiltrated leaves. For B and D, values marked with lowercase letters are statistically significantly different from those for other groups marked with different letters. (P < 0.05, ANOVA with the Tukey-Kramer HSD test).

### Mutation in HPR1, another components of THO/TREX complex can also restore *pho1-7* **shoot growth but not mutations affecting tasiRNA biogenesis**

To investigate if TEX1 exerts its function in the restoration of *PHO1* expression via the THO/TREX complex, we crossed *pho1-7* to *hpr1-6* which is a mutant in another component of THO/TREX core complex [14]. Double mutant *pho1-7 hpr1-6* partially restored shoot growth while maintaining relatively low Pi contents comparable to the *pho1-7 tex1-6* but slightly higher than *pho1-7* (Fig 1B and Fig 4 A-B).

**Fig 4.**
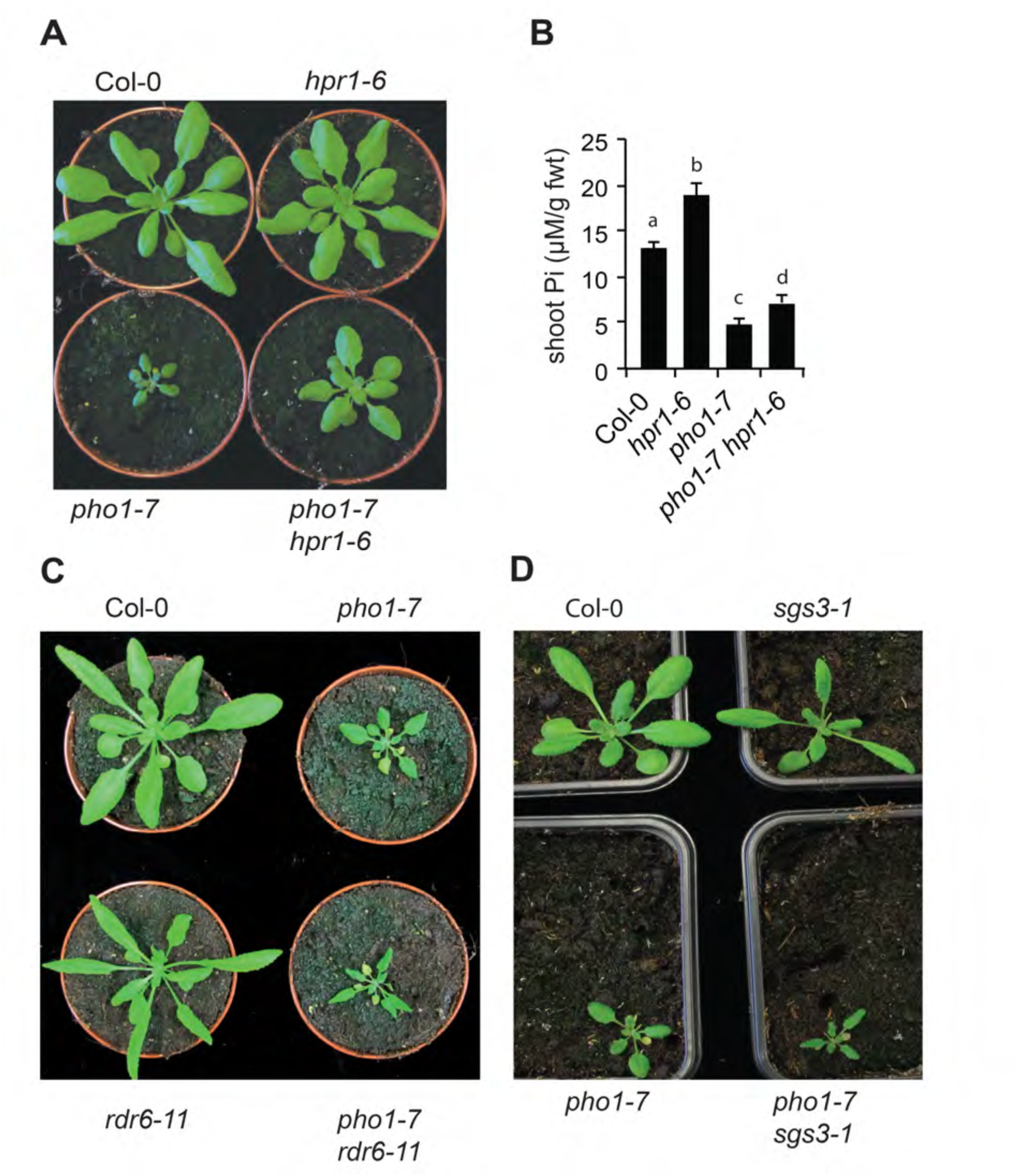
*hpr1* can restore *pho1-7* shoot growth but not mutations in tasiRNA biogenesis. (A, C, D) Phenotype of four-week-old plants. (B) Pi contents in the rosettes of four-week-old plants. Data from a representative experiment shows the means of Pi contents from ten different plants grown in independent pots. Errors bars represent standard deviation. Values marked with lowercase letters are statistically significantly different from those for other groups marked with different letters (P < 0.05, ANOVA with the Tukey-Kramer HSD test).

THO/TREX complex has previously been demonstrated to participate in the biogenesis of trans acting small interfering RNAs (tasiRNAs) and other small RNAs (siRNAs and miRNAs) that can affect levels of transcription through DNA methylation and some unknown mechanisms. We explored the possibility that changes in the biogenesis of tasiRNAs may be responsible for the growth phenotype associated with the *pho1-7 tex1-4* and *pho1-7 hpr1-6* mutants. We crossed *pho1-7* with two mutants in genes encoding core components required for the biogenesis of tasiRNAs, namely *rdr6-11* and *sgs3-1* [28, 29]. Double mutants *pho1-7 rdr6-11* and *pho1-7 sgs3-1* had shoot growth similar to the parental *pho1-7* (Fig 4C-D). Together, these results indicate disruption in distinct genes of the THO/TREX complex, namely *TEX1* and *HPR6*, can revert the growth phenotype of the *pho1-7* mutant and that biogenesis of tasiRNAs is not implicated in these processes.

### Impaired mRNA termination at the NOS terminator restores expression of truncated PHO1 in *pho1-7 tex1-4* mutant

To elucidate how mutations in *TEX1* and *HPR1* lead to changes in transcription at the *PHO1* locus, we performed paired-end next generation RNA sequencing of Col-0, *pho1-7*, *pho1-7 tex1-4* and *pho1-7 hpr1-6* mutants from roots. We first mapped the RNA reads of Col-0 and *pho1-7* against the Col-0 genome and confirmed the absence of the second exon of *PHO1* in the genome of *pho1-7* (S3A Fig). To understand the transcription dynamics at the *PHO1* locus in the various mutants derived from *pho1-7*, we mapped RNA sequencing reads to the *pho1-7* genomic configuration with the T-DNA insertion and exon 2 deletion. Detailed analysis of mRNAs from *PHO1* locus in *pho1-7* mutants indicated that transcription was initiated in the *PHO1* promoter and terminated at two different locations, namely at *NOS* terminator inside the T-DNA and at the endogenous *PHO1* transcription termination site (Fig 5A-C). Using RT-PCR and various primer combinations, we could detect four types of transcripts in the *pho1-7* mutant, namely one unspliced and two spliced mRNA ending at the *NOS* terminator, and one long transcript ending at the endogenous *PHO1* terminator and where *PHO1* exons 1 and 3 were appropriately spliced, removing the T-DNA and generating the *PHO1*^Δ249-342^ RNA variant (Figure 5A-D). While in the *pho1-7* mutant the four transcripts were expressed at similar low level, in *pho1-7 tex1-4* and *pho1-7 hpr1-6* double mutants the majority of transcripts was the *PHO1*^Δ249-342^ RNA variant (Fig 5C-D). Analysis by PacBio sequencing of full-length mRNAs produced at the *PHO1* locus in Col-0, *pho1-7*, *pho1-7 tex1-4* and *pho1-7 hpr1-6* supported to these conclusions and highlighted that essentially two classes of transcripts are produced in the various mutants, namely transcripts that include the 5’portion of the T-DNA and end at the NOS terminator and transcripts that end at the *PHO1* terminator and exclude the complete T-DNA (S3B Fig). While in the *pho1-7* mutant the majority of transcripts were of the first type, the *pho1-7 tex1-4* and *pho1-7 hpr1-6* mutants mostly expressed the second type. Such pattern of transcripts are not consistent with alternative splicing but rather indicate that transcription termination at the *NOS* terminator was suppressed in *pho1-7 tex1-4* and *pho1-7 hpr1-6* mutants, and this enabled the transcription machinery to reach the *PHO1* terminator and generate a transcript where exon 1 was spliced to exon 3, resulting in the removal of the T-DNA. Chromatin immunoprecipitation using an antibody against the elongating RNAPII (phosphorylated at S2 of the C-terminal domain) followed by qPCR showed that RNAPII occupation at the *PHO1* locus situated after the T-DNA insertion was significantly reduced in *pho1-7* mutants but increased in *pho1-7 tex1-4* mutant (Fig 5E), consistent with an increase in transcriptional read-through past the T-DNA in the *pho1-7 tex1-4* double mutant (Fig 5D).

**Fig 5.**
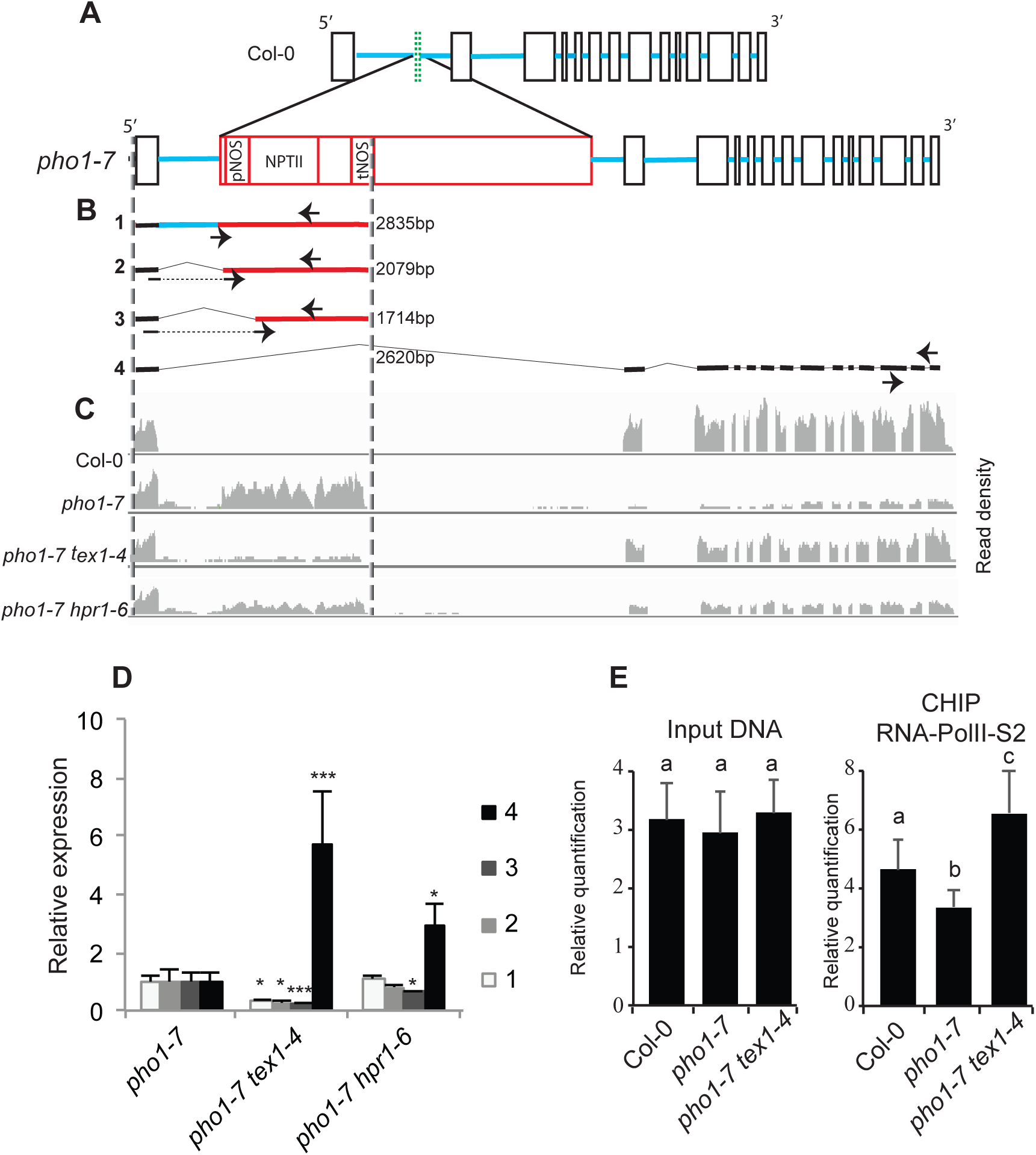
*tex1* mutant impairs transcription termination at the NOS terminator. (A) Structure of the wild type *PHO1* gene (top) and T-DNA (red) integrated in the *pho1-7* mutant (bottom). Exons are indicated as black open boxes, except for the second exon, indicated as a green doted box. Exon 2 is present in Col-0 but deleted in the *pho1-7* mutant as a result of T-DNA insertion, which is shown in red. Blue lines represent introns. pNOS, NOS promoter; NPTII, neomycin phosphotransferase gene; tNOS, NOS terminator. (B) Structure of the four mRNA variants produced at the *PHO1* locus in *pho1-7* mutants. The thick black and blue lines represent sequences derives from *PHO1* exons and introns, respectively, while the thick red lines are derived from the T-DNA. The sizes of the different transcripts are shown. Arrows indicate the location of primers used for the qPCR shown in panel D. Forward primers (discontinued arrows) in transcript 2 and 3 are at the junction of cryptic splicing sites to ensure specificity. (C) Illumina RNA sequencing reads density graph showing mRNA expression at the *PHO1* locus in various genotypes. The RNA sequencing reads are mapped against the *PHO1* locus in *pho1-7,* represented in the lower section of panel A (D) Quantification via qPCR of four different mRNA transcripts produced at the *PHO1* locus in the roots of 4-week-old plants. The numbers associated with each transcript and primers used for qPCR are shown in panel B. (E) CHIP-qPCR experiments showing the density of RNAPII-S2 at the 3’end of the *PHO1* gene. For D and E, data are means of three samples from plants grown in independent pots and three technical replicates for each sample. Error bars represent standard deviation. For D, statistical analysis was performed comparing each *PHO1* transcript isoform in *pho1-7 tex1-4* and *pho1-7 hpr1-6* double mutants relative to the *pho1-7* parent, with asterisks denoting statistical significance (*, P < 0.05; and ***, P < 0.001) according to Student’s t test. For E, values marked with lowercase letters are statistically significantly different from those for other groups marked with different letters (P < 0.05, ANOVA with the Tukey-Kramer HSD test).

### THO/TREX complex is required for the correct termination of mRNAs in endogenous loci

To understand if TEX1 and HPR1 contribute to the termination of RNA transcription at endogenous genes, RNA sequencing data generated from roots of Col-0, *pho1-7*, *pho1-7 tex1-4* and *pho1-7 hpr1-6* grown in soil for 3 weeks were analyzed for the presence of 3’ UTR extensions. We observed significant changes in 3’UTR extensions in *pho1-7 tex1-4* and *pho1-7 hpr1-6* mutants as compared to both Col-0 and *pho1-7.* Two examples of such 3’UTR extensions are shown in Fig 6A. While 3’UTR extensions were observed in only 3 transcripts in *pho1-7*, 72 and 51 transcripts showed 3’UTR extensions in the *pho1-7 tex1-4* and *pho1-7 hpr1-6* mutants, respectively, but not in the *pho1-7* parent, with a subset of 38 transcripts found in common between *pho1-7 tex1-4* and *pho1-7 hpr1-6* but not *pho1-7* (Fig 6B). These results indicate that while a large proportion of genes affected in their 3’UTR extension in the *tex1-4* respond similarly in the *hpr1-6*, the effects of these two mutations on RNA termination are not completely redundant.

**Fig 6.**
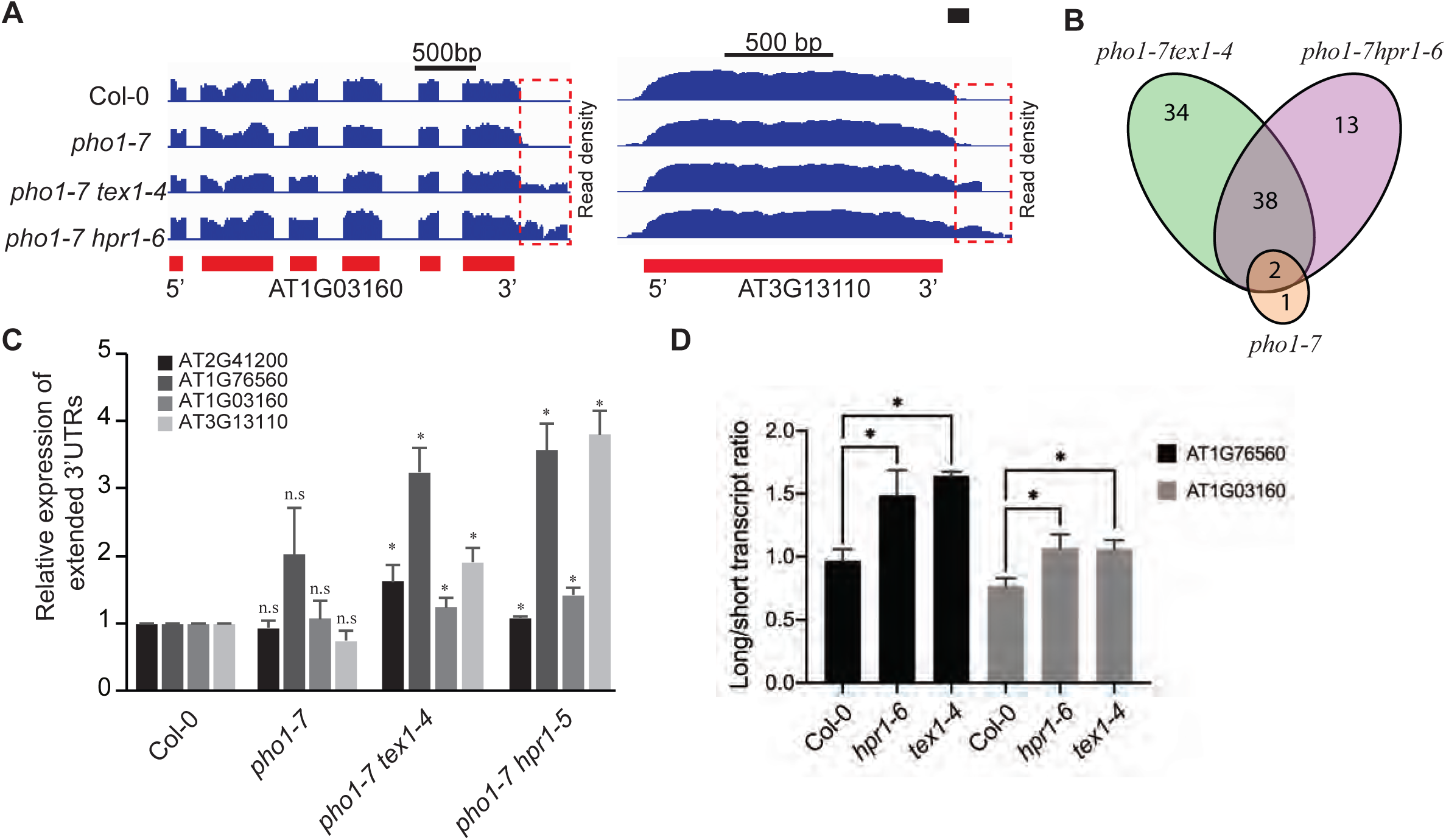
3′ UTR extensions in endogenous genes of the *pho1-7 tex1-4* and *pho1-7 hpr1-6* mutants. (A) Illumina RNAseq reads density maps (blue) showing examples of 3’ UTR extensions for genes AT1G03160 (left) and AT3G11310 (right) in the *pho1-7 tex1-4* and *pho1-7 hpr1-6* mutants. The 3′ UTR extensions are depicted by a red rectangle. The exons of the genes are indicated as red boxes below. (B) Venn diagram showing the number and overlap in genes with 3’UTR extensions. (C) Validation of 3’ UTR extensions via qPCR. (D) Transient expression of Luciferase gene fused after the stop codon to the 3’end of genes AT1G76560 and AT1G03160. The constructs were expressed in Arabidopsis mesophyll protoplasts obtained from Col-0, *hpr1-6* and *tex1-4*. The bar chart shows the relative ratios of long-to-short transcripts in the various genotypes. For C, data are means of three samples with each sample being a pool of seven seedlings and three technical replicates for each sample. For D, data are means of four samples with each sample being an independent transfection of protoplasts. Error bars represent standard deviation. Asterisks denote statistical significance (P < 0.05) from the Col-0 control according to Student’s t test (n.s.= not statistically different).

We hypothesized that if regulation of RNA termination by THO/TREX complex is generic and robust, changes in 3’UTR extensions should be relatively insensitive to growth conditions. To assess the robustness of 3’UTR extensions, we analyzed an independent RNA sequencing dataset generated from roots of Col-0 and *tex1-4* mutant grown *in vitro* for 7 days in MS medium supplemented with sucrose. A total of 77 transcripts showed 3’UTR extensions in the *tex1-4* mutant relative to Col-0, with 48 transcripts found also in the dataset of 3’UTR extensions for *pho1-7 tex1-4* mutant grown for 3 weeks in soil (S4 Fig), indicating that a large proportion of transcripts with 3’UTR extensions observed in the *tex1-4* genetic background are insensitive to major changes in growth conditions.

Validation of 3’UTR extensions in a set of genes identified by RNA sequencing was first performed by qPCR using (Fig 6C). A transient assay was also developed whereby the sequence 500 bp downstream of the stop codon (which includes the 3’UTR) of two genes, AT1G76560 and AT1G03160, was fused after the stop codon of the nano-luciferase gene. These constructs were expressed in Arabidopsis mesophyll protoplasts and the ratio of transcripts with an extended 3’end relative to the main transcription termination site was determined by qRT-PCR 16 hours after transformation. Analysis revealed an increase, for both constructs, in the ratio of long-to-short transcripts by approximately 50-60% in the *hpr1-6* and *tex1-4* mutants compared to Col-0 (Fig 6D), further supporting the implication of both TEX1 and HPR1 in mRNA termination.

GO term enrichment analysis of transcripts with impaired RNA termination in the *tex1-4, pho1-7 tex1-4* and *pho1-7 hpr1-6* showed that these transcripts did not belong to a particular functional category (S5 Fig). Therefore, we looked at the sequences of RNA termination sites for mechanistic clues of impaired RNA termination and 3’UTR extension in *tex1* and *hpr1* mutants. RNA termination sites are defined characteristic motifs, including an A-rich region at approximately −20 nucleotides (position −1 being defined as the last nucleotide before the polyA tail), which includes the AAUAAA-like sequence. This is followed by a U-rich region at −7 nucleotide and a second peak of U-rich region at approximately +25 nucleotides [30–32]. Analysis of the distribution of nucleotides upstream and downstream of the 3’ cleavage site did not reveal a significant difference from this distribution for genes showing a 3’extension in the *tex1-4* and *hpr1-6* mutant backgrounds (Fig 7A). Additionally, we analyzed the differences at −20 polyadenylation signal for the genes with 3’ extensions compared to all Arabidopsis genes. Although not statistically significant (p=0.22), a lower representation of the canonical AAUAAA sequence appeared associated with the group of genes with 3’ extension compared to all genes (Fig 7B). We used the transient expression to test the effect of changing the single AAUGAA polyadenylation signal present in gene AT1G76560 to the canonical AAUAAA. While the optimized polyadenylation signal led to a small decrease in 3’UTR extensions relative to the wild type sequence in Col-0, there was still an approximately 50% increase in the ratio of long-to-short transcripts in the *hpr1-6* and *tex1-4* mutant compared to Col-0 (Fig 7C). Altogether, these results indicate that while HPR1 and TEX4 contribute to mRNA termination, they do not appear to act primarily via the nature of the −20 polyadenylation signal.

**Fig 7.**
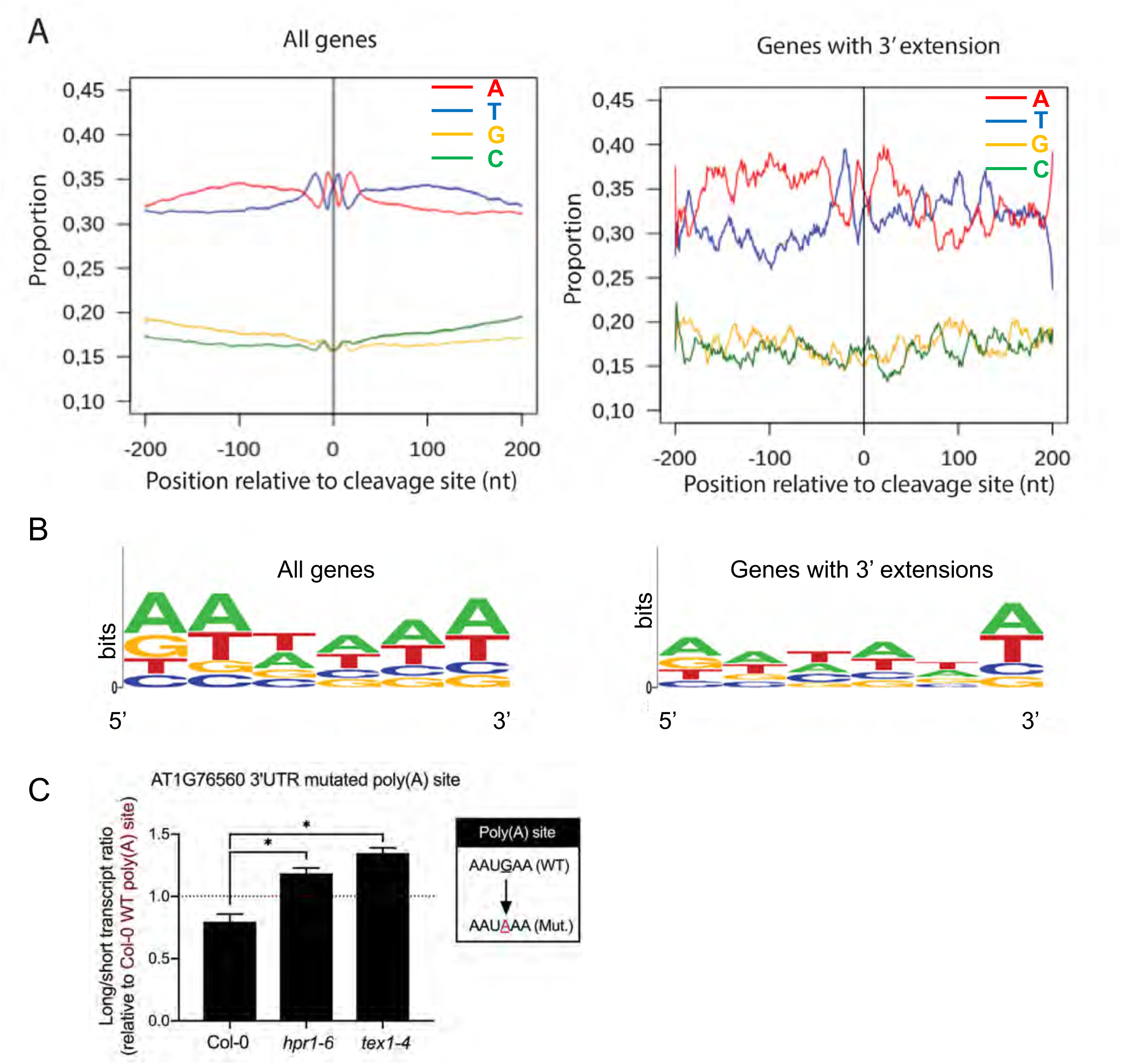
Nucleotide composition near the 3’ cleavage site of mRNAs. (A) Proportion of bases in a 200 nucleotide region 5’ and 3’ of the transcripts termination sites for all genes (left) or genes showing 3’UTR extensions in the *hpr1-6* and *tex1-4* mutant background (right). (B) Graphical representation of base enrichments found at the near upstream element sequence of all genes (left) and genes with 3’UTR extensions in the *hpr1-6* and *tex1-4* mutant background (right). Chi square test pvalue = 0.22 (C) Transient expression of Luciferase gene fused after the stop codon to the 3’end of the gene AT1G76560 in which the wild-type polyadenylation site AAUGAA was mutated to AAUAAA. The construct was expressed in Arabidopsis mesophyll protoplasts obtained from Col-0, *hpr1-6* and *tex1-4*. The bar chart shows the ratios of long-to-short transcripts in the various genotypes and values are expressed relative to the wild-type construct in Col-0. Data are means of four samples with each sample being an independent transfection of protoplasts. Error bars represent standard deviation. Asterisks denote statistical significance (P < 0.05) from the Col-0 control according to Student’s t test.

## Discussion

The contribution of the THO/TREX complex to mRNA biogenesis has been particularly studied for splicing and export [9]. In contrast, the direct implication of the THO/TREX complex in mRNA termination is poorly defined and reported only in few studies in human [33, 34]. The THO5 was shown to interact with both CPSF100 and CFIm, two proteins involved in 3’end-processing and polyadenylation site choice, and differences in mRNA polyadenylation were observed in human cells depleted for THO5 [33, 34]. Recruitment of the cyclin-dependent kinase CDK11 by the THO/TREX complex was shown to be essential for the phosphorylation of RNAPII and the proper 3’end processing of the human immunodeficiency virus RNA, although it is unknown if this interaction is also mediated via THO5 [35].

Although considerably less is known about mRNA biogenesis in plants compared to yeast and metazoans, proteins forming the THO core complex are also found in plants, implicating a conservation in their mode of action [13]. Indeed, *A. thaliana* mutants in the *HPR1* and *TEX1* genes show defects in mRNA splicing and export [18, 19, 36]. However, most of our knowledge in plants on THO core complex relates to its implication in small RNA biogenesis. *A. thaliana* mutants in either *HPR1*, *TEX1*, *THO2* or *THO6* are defective in the synthesis of one or multiple forms of small RNAs, including miRNA, siRNA, and tasiRNAs, although the mode of action behind this defect is currently unknown [14–17]. Some of the phenotypes associated with the *tex1* and *hpr1* mutants in *A. thaliana*, such as repression of female germline specification or reduction in scopolin biosynthesis under abiotic stress, are likely caused by a reduction in tasiRNA or miRNA production [37, 38].

This current work highlights the contribution of both TEX1 and HPR1 to mRNA termination. Mutations in these genes in a *pho1-7* mutant background having a T-DNA insertion between the *PHO1* exons 1 and 3 led to the suppression of mRNA termination at the NOS terminator present in the T-DNA, followed by transcriptional read-through and splicing of the T-DNA, resulting in the generation of a near full-length *PHO1* mRNA (minus exon 2). The truncated PHO1 protein generated from this mRNA maintained some Pi export activity, resulting in a reversion of the growth phenotype associated with the severe Pi deficiency of the *pho1-7* mutant.

Beyond its effect on the NOS terminator, mutations in the *HPR1* and *TEX1* genes also affected mRNA 3’ processing of endogenous loci leading to 3’ UTR extensions. Interestingly, the majority of loci with impaired transcription termination were shared between *tex1* and *hpr1* mutants, suggesting that both proteins have overlapping functions in RNA termination. Analysis of nucleotides surrounding of the 3’ cleavage site did not reveal a significant difference for genes showing a 3’extension in the *tex1-4* and *hpr1-6* mutants compared to the Arabidopsis genome. The best defined DNA sequence involved in the control of mRNA polyadenylation site is a A-rich region at approximately −20 nucleotides defined as the near upstream element (NUE). Although the NUE canonical AAUAAA sequence is found very frequently in animal genomes, the heterogeneity in NUE sequences is larger in plants [30, 39]. An apparent deviation (but not statistically significant) from the AAUAAA was observed in genes showing a 3’UTR extension in the *hpr1-6* and *tex1-4*. Furthermore, optimization of the polyadenylation site of the gene AT1G76560 from AAGAAA to AAUAAA did not affect the extent of 3’UTR extension in the *tex1-4* and *hpr1-6* mutants compared to Col-0. It is likely that the relatively low number of genes showing 3’UTR extensions significantly limits our ability to identify nucleotide features that are important in 3’UTR extension in the *tex1-4* and *hpr1-6* mutants through a bioinformatic approach. A more systematic analysis of the 3’UTR of the genes affected in the *tex1-4* and *hpr1-6* mutants, such as AT1G76560 and AT1G03160, using the transient assay described in this study may lead to the identification of the cis-elements involved.

Analysis of the *A. tumefaciens* NOS transcript revealed two putative NUE sequences, namely AAUAAA and AAUAAU, at position −135 and −50 nucleotides, which is much further upstream than the usual −20 nucleotides [40]. It is thus likely that other sequences within the NOS terminator play a determinant role in RNA transcription termination. Furthermore, while the prominent dinucleotide located immediately before the cleavage site are typically CA or UA, the dinucleotide GA is present in the NOS terminator [40]. While no detailed functional analysis of the polyadenylation signal of the NOS gene has been reported, it is likely that the its unusual structure is related to the bacterial origin of the gene. While the NOS terminator is a common feature of many T-DNA vectors, several studies have shown that transgene expression can be considerably enhanced either when the NOS terminator is combined with a second terminator or when it is replaced by the terminator of plant endogenous genes [41–43]. For example, replacement of the NOS terminator with an extensin terminator was shown to reduce read-through transcript and improve expression of transgenes [44]. Altogether, these data suggest that mutations in the *HPR1* and *TEX1* genes may more prominently affect mRNA 3’processing for genes having unusual or weak polyadenylation signals, such as found in the NOS terminator.

The 3’UTR of mRNAs have important regulatory functions, impacting mRNA stability, localization and translation potential via interaction with numerous RNA binding proteins as well as miRNAs [45]. Extension of the 3’UTRs of genes in the *hpr1-6* and *tex1-4* genetic background may thus impact their expression in numerous ways. In some cases, extension of 3’UTR may also lead to disruption of the downstream gene by the formation of an antisense RNA with potential to trigger siRNA-mediated gene silencing, or by transcriptional interference via RNAPII collision [46]. The *tex1* and *hpr1* mutant are known to have multiple phenotypes, ranging from defects in vegetative and reproductive development [18, 38], responses to both biotic and abiotic stress [36, 37] and the expression of genes encoding acid phosphatases [16] or ethylene signaling pathway repressor [20]. It would be of interest to determine if the genes affected by 3’UTR extensions contribute to some extent to these phenotypes.

Further work is necessary to gain detailed mechanistic insights as to how mutations in *tex1* and *hpr1* lead to both suppression of termination at the NOS terminator and changes in the 3’UTR of endogenous genes. Being part of the TREX complex associated with the mRNA transcription machinery, TEX1 and HPR1 could affect mRNA transcription termination through interactions with proteins more specifically involved in 3’end processing. This would be analogous to the implication of the THOC5 protein, another component of the TREX complex, in mRNA 3’-end processing in mammals via interactions with mRNA cleavage factors, including CPSF100 [33, 34]. Since TEX1 and HPR6 have both been implicated in the generation of small RNAs, including tasiRNA, siRNA and miRNA, the contribution of these pathway to mRNA termination should also be further examined. Reversion of the *pho1-7* growth phenotype could not be reproduced by mutations in the *SGS3* and *RDR6* genes involved in small RNA biogenesis, in particular of tasiRNAs, indicating that the effects of the *hpr1* and *tex1* mutations in *pho1-7* could not be caused by changes in tasiRNAs generation [14, 15, 17, 18]. siRNA is associated with DNA methylation, which in turn could impact RNA polymerase activity and mRNA processing, including splicing and termination [47, 48]. Although the fact that T-DNA present in the *pho1-7* mutant is both well transcribed and mediates resistance to kanamycin, suggesting that it is unlikely to be strongly methylated, more subtle effects of siRNA-mediated methylation on RNA transcription termination at the NOS terminator and endogenous loci should be explored.

The N-terminal half of the PHO1 protein harbors a SPX domain which is involved in binding inositol polyphosphate at high affinity via interactions with conserved tyrosine and lysine residues [49, 50]. A PHO1 protein with mutations in these SPX conserved amino acids is unable to complement the *pho1-2* null mutant, indicating that the binding of inositol polyphosphate is important for PHO1 activity *in plantae* [50]. The protein PHO1^Δ84-114^ synthesized from the *pho1-7* mutant does not affect the core of the SPX domain but only leads to a small deletion at the N-terminal end of the second SPX subdomain (see S1 text). Heterologous expression of the PHO1^Δ84-114^ protein in tobacco leaves led to specific Pi export activity, indicating that the 31 amino acid deletion does not completely inactivates the protein. Considering that *pho1-7 tex1-4* and *pho1-7 hpr1-6* as well as the *pho1-4* null mutant expressing the PHO1^Δ84-114^ protein have low shoot Pi, is thus likely that the PHO1^Δ84-114^ retains some Pi export activity in root xylem parenchyma cells, but lower than the wild type protein. The high level of expression of the PHO1^Δ84-114^ protein in the *pho1-7 tex1-4* mutant cannot be simply explained by a higher expression of the truncated *PHO1*^Δ249-342^ mRNA in the double mutant background relative the *pho1-7* parent, since the *PHO1*^Δ249-342^ mRNA remained lower than the full length *PHO1* mRNA in Col-0 plants. PHO1 is known to be degraded by PHO2, a key protein involved in the Pi-deficiency signaling pathway [51]. Whether the high level of PHO1^Δ84-114^ accumulation is a reflection of greater stability of the truncated protein and/or increased translation efficiency of the truncated mRNA remains to be determined.

Uncoupling low leaf Pi from its main effect on shoot growth has previously been reported for plants under-expressing *PHO1* via silencing, indicating a role for high root Pi content and PHO1 in modulating the response of the shoot to Pi deficiency [25]. The improved shoot growth observed in the *pho1-7 tex1-4* and *pho1-7 hpr1-6* compared to the parent *pho1-7* is likely due to the increased expression of the PHO1^Δ84-114^ hypomorphic protein and an increase in Pi export activity. However, both *hpr1* and *tex1* mutants have low expression of the *RTE1* gene involved in the repression of the ethylene signaling pathway which has been linked to an increase in root-associated acid phosphatase activity and root hair elongation in these mutants, two characteristics that can positively impact Pi acquisition [16, 20]. It is thus possible that a small part of the growth improvement observed in the *tex1-4 pho1-7* and *hpr1-6 pho1-7* may also be associated with the repression of the ethylene pathway.

## Materials and methods

### Plant material and growth conditions

All Arabidopsis plants used in this study, including mutants and transgenic plants, were in the Columbia (Col-0) background. For *in vitro* experiments, plants were grown in half-strength Murashige and Skoog (MS) salts (Duchefa M0255) containing 1% sucrose and 0.7% agar. For Pi-deficient medium MS salts without Pi (Caisson Labs, MSP11) and purified agar (Conda, 1806) was used. Pi buffer pH5.7 (93.5% KH_2_PO_4_ and 6.5% K_2_HPO_4_) was added to obtain different Pi concentrations. In the Pi-deficient media, Pi buffer was replaced by equimolar amounts of KCl. Plants were also grown either in soil or in a clay-based substrate (Seramis) supplemented with half-strength MS for the isolation of roots. Growth chamber conditions were 22°C, 60% humidity, and a 16h light/8h dark photoperiod with 100 µE/m2 per sec of white light for long days and 10h light/14h dark for short days. The *pho1-2*, *pho1-4*, *pho1-7* (SALK_119529) and *tex1-4* were previously described [15, 22, 52] and *hpr1-6* (SAIL_1209_F10) is a T-DNA mutant from SAIL collection. The *rdr6-11* is an EMS-derived mutant while *sgs3-1* is a T-DNA mutant from the SALK collection (SALK_001394) and have both been previously described [28].

### *pho1* suppressor screen

Ethyl methanesulfonate (EMS) mutagenesis was performed on approximately 20,000 seeds of homozygous *pho1-7*. Seeds were treated with 0.2% EMS in 100mM phosphate buffer for 8 hours and were rinsed with water 10 times afterwards. Approximately 10, 000 individual M1 plants were grown and seeds of their progeny were collected in bulk. Approximately 10’300 M2 plants were grown in soil for 4 weeks under an 18h/8h day/night light cycle. Plants showing an increased rosettes size relative to the *pho1-7* parent were identified and their seeds collected. In the next generation, plants retaining 100% kanamycin resistance mediated by the T-DNA in *PHO1* were sown in soil and the Pi content in 3-week-old rosettes was determined using the molybdate assay [53] as previously described [27]. Plants showing the combination of increased rosettes size with low shoot Pi similar to *pho1-7* were then genotyped by PCR to further confirm the presence of the T-DNA in the *PHO1* locus in an homozygous state.

### Identification of mutant genes by next generation sequencing

*pho1-7* suppressor mutant was back-crossed to the parent *pho1-7* line to test if the mutation was dominant or recessive and to generate an isogenic mapping population. Approximately 40 plants with a *pho1-7* suppressor phenotype were selected from the segregating F2 population. DNA was extracted from the pool of these 40 mutants and sequencing was performed using Illumina Hiseq 2000 system. DNA sequencing yielded an average coverage of 80 to 100 nucleotides per nucleotide per sample. Sequencing reads were mapped with Burrows-Wheeler Aligner software (BWA) version 0.5.9-r16 using default parameters to the TAIR10 release of the *A. thaliana* genome. Using SAM (Sequence Alignment/Map) tool the alignments were converted to BAM format. SNPs were called with the Unified Genotyper tool of the Genome Analysis Toolkit (GATK) version v1.4-24-g6ec686b. SNPs present in the parental line *pho1-7* were filtered out using BEDTools utilities version v2.14.2. TAIR10_GFF3_genes_transposons.gff file was used to filter out the SNPs present in the transposons. The predicted effect of the remaining SNPs in coding regions was assessed with SNPEff version 2.0.4 RC1. Unix command awk was used to extract the SNP frequencies (the number of reads supporting a given SNP over the total number of reads covering the SNP location) and were plotted with R 2.15.1.

### Cloning and transgenic lines

For complementation of *pho1-7 suppressor* with *TEX1*, genomic sequence including 2 kb promoter 5’UTR and 3’UTR was amplified using primers TEX1-gen-F -and TEX1-gen-R (sequences of all primers used are listed in S1 Table). The amplicon was cloned into pENTR/D TOPO vector (Invitrogen). The entry vector was then shuttled into the binary vector pMDC99 [54] using Gateway technology (Invitrogen). For pTEX1:TEX1:GFP fusion, *TEX1* promoter and gene was amplified using primers TEX1-gen-F and TEX1-gen-R-w/o-stop. Reverse primer was designed to remove the stop codon from *TEX1* gene. The amplicon was cloned into pENTR/D TOPO vector (Invitrogen). The entry vector was then shuttled into the binary vector pMDC107, which contains GFP at the C-terminal [54]. For TEX1 promoter GUS fusion, promoter was amplified using the primers TEX1-Pro-infu-LP and TEX1-Pro-infu-RP. The amplicon was cloned into the gateway entry vector pE2B using Infusion technology (Clonetech). The entry vector was then shuttled into the binary vector pMDC63 [54] using Gateway technology (Invitrogen). For cloning pPHO1:gPHO1^Δ83-114^:GFP, promoter and first exon was amplified using primers P2BJ pPHO1 1exon L and P2BJ pPHO1 1exon R (fragment 1), *PHO1* gene from third exon until before the stop codon was amplified using primers P2BJ PHO1gene 3rd exon F and P2BJ PHO1gene 3rd exon R (fragment 2). The two fragments were combined together in the Gateway entry vector pE2B using Infusion technology (Clonetech). The entry vector was then shuttled into the binary vector pMDC107 [54] using Gateway technology (Invitrogen). All the binary vectors were transformed into *Agrobacterium tumefaciens* strain GV3101 and plants were transformed using flower dip method [55].

For analysis of transcript termination by transient expression in Arabidopsis protoplasts, the 500 nucleotides located immediately after the stop codon of the genes AT1G76560 and AT1G03160 were fused after the stop codon of the nano-luciferase (nLUC) gene. To achieve this, the 500 bp 3’sequences from AT1G76560 (wild-type and mutated) and AT1G03160 flanked by attR1 and attR2 sites were synthesized and inserted in the pUC57 plasmid by Genscript. The DNA insert was then shuttled by Gateway cloning into the dual-luciferase vector nLucFlucGW (Genbank MH552885) [56] modified to lack the original nLuc 3’UTR and terminator sequences. The final constructs had the hybrid genes expressed under the control of the ubiquitin promoter, as well as the firefly luciferase gene (Fluc) constitutively expressed, used for loading control.

### Quantitative RT PCR and phosphate measurement and Pi export assay

Quantitative RT-PCR was performed as previously described [27]. For transient expression of PHO1, *Nicotiana benthamiana* tobacco plants were infiltrated with *A. tumefaciens* as previously described [21]. Pi measurements were performed using the molybdate-ascorbic acid method [53].

### Confocal microscopy

Seedlings were incubated for ten min in a solution of 15 mM Propidium Iodide (sigma, P4170) and rinsed twice with water. Excitation and detection window for GFP was set at 488 nm for excitation and 490–555 nm for detection. Propidium iodide was excited at 555 nm and detected at 600-700nm. All experiments were performed using Zeiss LSM 700 confocal microscope.

### Western blot analysis

Plants were grown on clay-based substrate (Seramis) supplemented with half-strength MS liquid medium. Proteins were extracted from homogenized 25-day-old roots at 4 °C in extraction buffer containing 10 mM phosphate buffer pH 7.4, 300 mM sucrose, 150 mM NaCl, 5 mM EDTA, 5 mM EGTA, 1 mM DTT, 20 mM NaF and 1× protease inhibitor (Roche EDTA free complete mini tablet), and sonicated for 10 min in an ice-cold water bath. Fifty micrograms of protein were separated on an SDS-PAGE and transferred to an Amersham Hybond-P PVDF membrane (GE healthcare). The rabbit polyclonal antibody to PHO1 [51] and goat anti-rabbit IgG-HRP (Santa Cruz Biotechnology, USA) was used along with the Western Bright Sirius HRP substrate (Advansta, USA). Signal intensity was measured using a GE healthcare ImageQuant RT ECL Imager.

### Illumina RNA-sequencing data analysis

RNA was extracted from roots of plants grown for 3 weeks in pots containing clay-based substrate (Seramis) or for 7 days on vertical agar plates containing half-strength MS media with 1% sucrose. Strand-specific libraries were prepared using the TruSeq Stranded Total RNA kit (Illumina). PolyA^+^ RNAs were selected according to manufacturer’s instructions and the cDNA libraries were sequenced on a HiSeq 2500 Illumina sequencer. The reads were mapped against TAIR10.31 reference genome using Hisat2 [57] and the readcount for each gene was determined using HTSeqcount [58]. Readcounts were normalized using DESeq2 [59]. Figures showing read density from RNAseq data were generated using Integrative genomics viewer (IGV) [60].

### Analysis of full-length mRNA using PacBio sequencing

One µg of total RNA was used to generate cDNA with the SMARTer PCR cDNA Synthesis kit (Clontech, Mountain View, CA, USA). Fifty µl of cDNA were amplified by 13 PCR cycles with the Kapa HiFi PCR kit (Kapa Biosystems, Wilmington, MA, USA) followed by size selection from 1.5kb to 3.5kb with a BluePippin system (Sage Science, Beverly, MA, USA). Seventy ng of the size selected fragment were further amplified with Kapa HiFi PCR kit for 5 cycles and 2 minutes extension time and 750 ng was used to prepare a SMRTbell library with the PacBio SMRTbell Template Prep Kit 1 (Pacific Biosciences, Menlo Park, CA, USA) according to the manufacturer’s recommendations. The resulting library was sequenced with P4/C2 chemistry and MagBeads on a PacBio RSII system (Pacific Biosciences, Menlo Park, CA, USA) at 240 min movie length using one SMRT cell v2. Bioinformatics analysis were performed through SMRT® Analysis Server v2.3.0. using RS_IsoSeq.1 workflow and TAIR10.31 as reference genome.

### Identification of 3’UTR extensions

3’ UTR extensions were identified following a procedure adapted from Sun et al. 2017 [8]. Briefly, reads obtained by single or paired-end polyA+ RNAseq were mapped with Hisat2 [57] against the intergenic regions extracted from TAIR10.31 annotation. Each intergenic region was divided into 10 nucleotide bins and the normalized readcount was determined for each bin with HTSeq-count [58] and DESeq2 [59]. 3’ extensions were then contiguously assembled from the 5’ end of intergenic intervals until a bin had a normalized readcount < 1. Only extensions longer than 200 nucleotides were kept for further analyses.

The number of reads mapping each TAIR10.31 gene and newly identified 3’ extensions was determined with HTSeq-count [58]. Differential expression analysis was performed with DESeq2 [59] to identify extensions significantly up or down-regulated independently of the expression level of the TAIR10.31 annotated gene body, comparing different genotypes. An extension was considered significantly differentially expressed if the adjusted pvalue corrected for false discovery rate was < 0.1 and the fold change of the ratios normalized readcount 3’ extension / normalized readcount gene body between 2 genotypes was > 2.

To analyze the polyadenylation signal present in genes with and without 3’UTR extensions, the frequency of each nucleotide at the polyadenylation consensus sequence AAUAAA was calculated for each gene and a Chi square test was used to test for statistical significance.

### Chromatin immunoprecipitation analyzed by qPCR

Leaves from 3-week-old *A. thaliana* seedlings from different genotypes were harvested and immediately incubated in 37 ml of pre-chilled fixation buffer (1% formaldehyde in 0.4M sucrose, 10 mM Tris pH 8, 1mM EDTA, 1 mM PMSF, 0.05% Triton X-100) for 10 min under vacuum. 2.5 ml of Glycine (2.5 M) was added and samples were incubated 5 additional min under vacuum, rinsed 3 times with water and frozen in liquid nitrogen. Frozen samples were ground and the powder resuspended in 30 ml of extraction buffer I (0.4 M sucrose, 10 mM HEPES pH 8, 5 mM ß-mercaptoethanol, 0.1 g/ml 4-(2-aminoethyl) benzenesulfonyl fluoride hydrochloride (AEBSF). After 20 min incubation at 4°C, the mixture was filtered through Miracloth and centrifuged for 20 min at 3000 g at 4°C. The pellet was resuspended in 300 µl of extraction buffer III (1.7 M sucrose, 10 mM HEPES pH 8, 0.15% Triton X-100, 2 mM MgCl_2_, 5 mM 5 mM ß-mercaptoethanol, 0.1 g/ml AEBSF), loaded on top of a layer of 300 µl of extraction buffer III and centrifuge for 1h at 16000 g at 4°C. The pellet was resuspended in 300 µl of Nuclei Lysis Buffer (50 mM HEPES pH 8, 10 mM EDTA, 1% SDS, 0.1 g/ml AEBSF) and incubated on ice for 30 min.

Chromatin solution was centrifuged twice for 10 min at 14000 g, 4°C and incubated over-night with S2P-RNApolII specific antibodies. The mixture was then incubated with Protein A beads for 3h at 4°C. After washing, immune complexes were eluted twice with 50 µl of Elution Buffer (1 % SDS, 0.1 NaHCO3). To reverse crosslinking, 4 µl of a 5 M NaCl solution was added to 100 µl of eluate and the mixture incubated overnight at 65 °C. Two µl of 0.5 M EDTA, 1.5 µl of 3 M Tris-HCl pH 6.8 and 20 µg of proteinase K were then added and the mixture incubated for 3h at 45 °C. DNA was then extracted using the NucleoSpin kit from Macherey Nagel. DNA samples were diluted 10 times and 2 µl were used for quantification by qPCR using Master Mix SYBR Select (Applied Biosystems).

### Transient expression in Arabidopsis protoplasts

Arabidopsis protoplasts were produced and transformed as previously described [61]. In brief, wild type Col-0, as well as *hrp1-6* and *tex1-4* mutant plants were grown in long photoperiod (16 h light and 8 h dark at 21 ^0^C) for 4-5 weeks and leaves were cut with razor blades to produce 0.5-1 mm leaf strips. These were submerged in enzyme solution (1% cellulase, 0.25% macerozyme, 0.4 M mannitol, 20 mM KCl, 20 mM MES and 10 mM CaCl2), vacuum infiltrated and incubated at room temperature for 2 h. Protoplasts were harvested by centrifugation at 100 xg for 3 min, washed with W5 solution (154 mM NaCl, 125 mM CaCl2, 5 mM KCl and 2 mM MES) and resuspended in MMG solution (4 mM MES, pH 5.7, 0.4 M mannitol and 15 mM MgCl2) at 1×10^6^ protoplast/ml. Protoplast transformation was performed by combining ∼1.5 x10^5^ protoplasts, 8µg of plasmid, and PEG solution (40% PEG4000, 0.2 M mannitol and 100 mM CaCl2). After replacing PEG solution with W5 solution by consecutive washings, protoplasts were kept in the dark for approximately 16 hours at 21^0^C. Transformed protoplasts were harvested by centrifugation at 6000 xg for 1 min, and resuspended in 1X Passive Lysis buffer (Promega, E1941). The lysate was cleared by centrifugation and RNA was extracted using RNA purification kit as described by the manufacture (Jena Bioscience, PP-210), followed by DNase I treatment. cDNA was synthesized from 0.1 µg RNA using M-MLV Reverse Transcriptase (Promega, M3681) and oligo d(T)15 as primer following the manufacturer’s instructions. qPCR analysis was performed using SYBR select Master Mix (Applied Biosystems, 4472908) with primer pairs specific to transcripts of interest and firefly luciferase mRNA, used for data normalization. Long/short transcript ratio was calculated with the following formula:

ΔCT(long transcript) = CT(long transcript) - CT(Fluc)

ΔCT(short transcript) = CT(short transcript) - CT(Fluc)

ΔΔCT(long/short) = ΔCT(long transcript) / ΔCT(short transcript) Long/short transcript ratio= 2^ΔΔCT(long/short)^.

## Acknowledgment

The authors are grateful to Syndie Delessert for technical assistance and to Tzyy-Jen Chiou (Academia Sinica, Taiwan) for the PHO1 antibody.

## Supporting Information Supporting Figure Legends

**S1 Fig.**
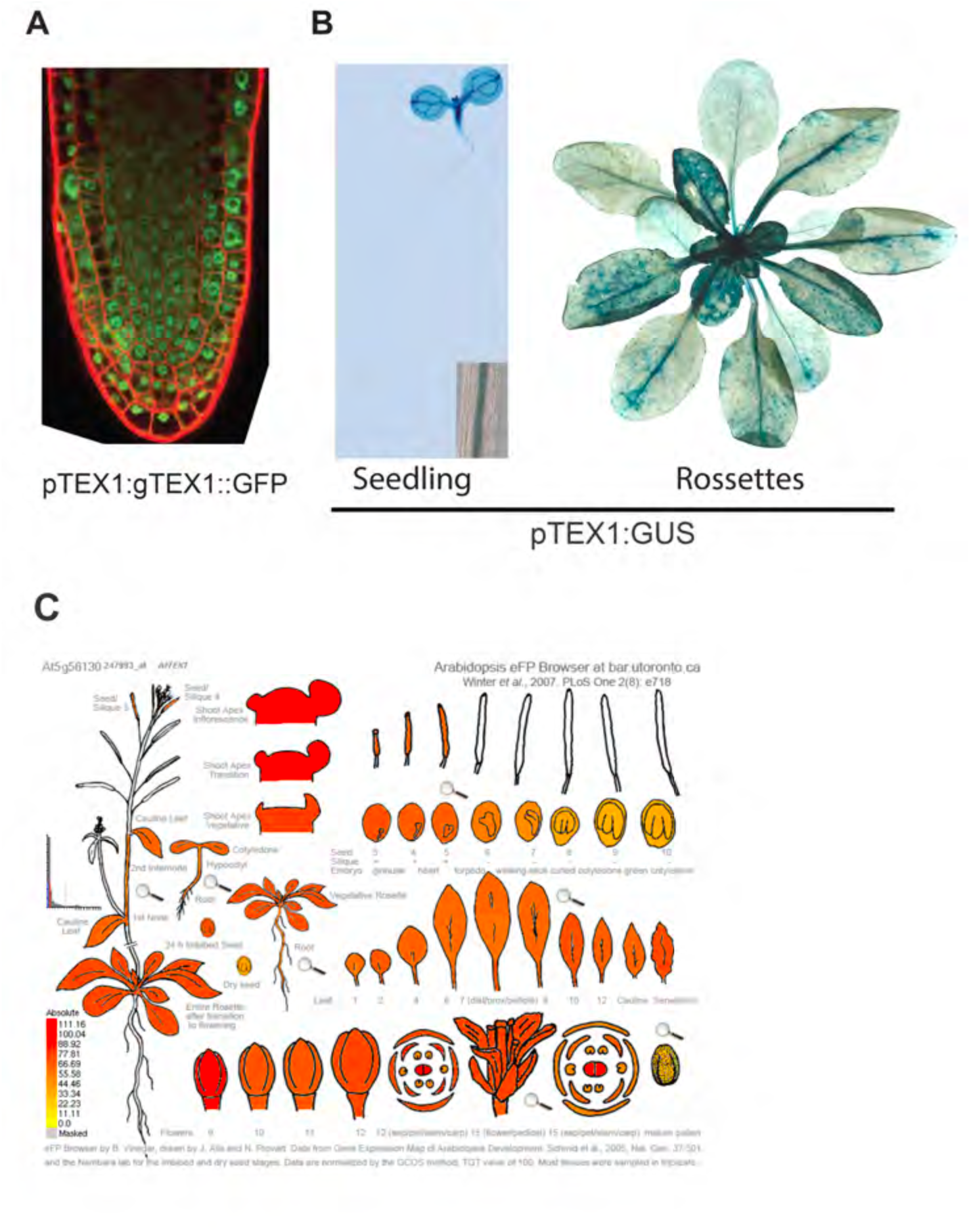
TEX1 is broadly expressed and localized in the nucleus. (A) TEX1 localization in root tips of 5-day-old seedlings. Expression of *TEX1::GFP* fusion is under the control of the endogenous *TEX1* promoter. (B) GUS expression from the *TEX1* promoter in roots and cotyledons of 5-day-old seedlings (left) as well as rosette leaves of 4-week-old plants (right). Plants were transformed with a *TEX1 promoter:GUS* construct. Inset (lower left) shows expression in a section of mature root. (C) Expression profile of the *TEX1* gene in Arabidopsis as visualized with the eFP Browser 2.0 (https://bar.utoronto.ca/efp2/).

**S2 Fig.**
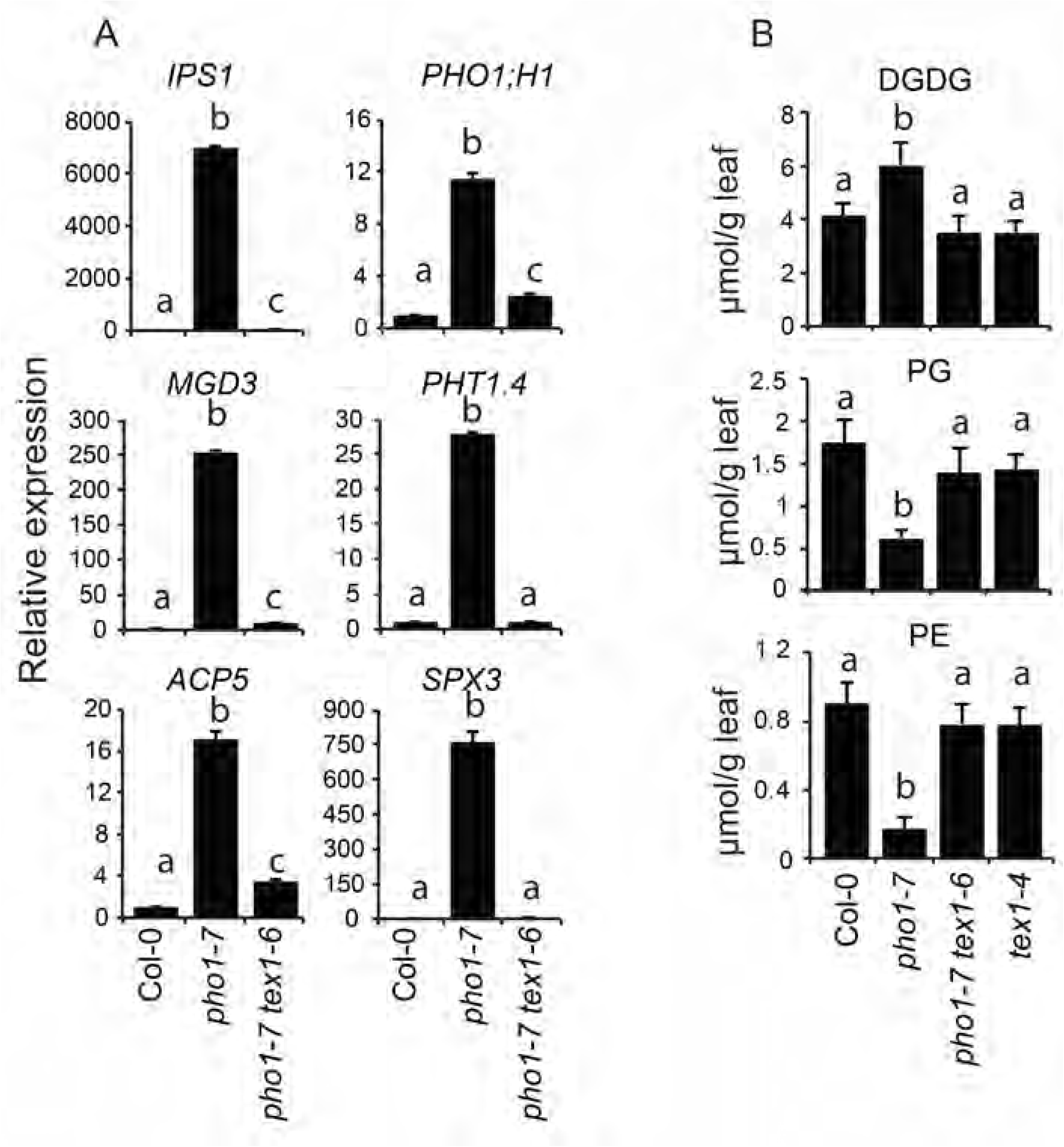
*pho1-7* suppressor mutants restores the expression of phosphate starvation response genes and lipid dynamics comparable to Col-0 level. (A) Relative expression of phosphate starvation-induced genes in the shoots of 4-week-old plants. (B) Lipid quantification in the shoots of 4-week-old plants. DGDG, digalactosyldiacylglycerol; PG, phosphatidylglycerol; PE, phosphatidylethalonamine. Data in A and B are means of three samples from plants grown in independent pots and three technical replicates. Error bars represent standard deviation. Values marked with lowercase letters are statistically significantly different from those for other groups marked with different letters (P < 0.05, ANOVA with the Tukey-Kramer HSD test).

**S3 Fig.**
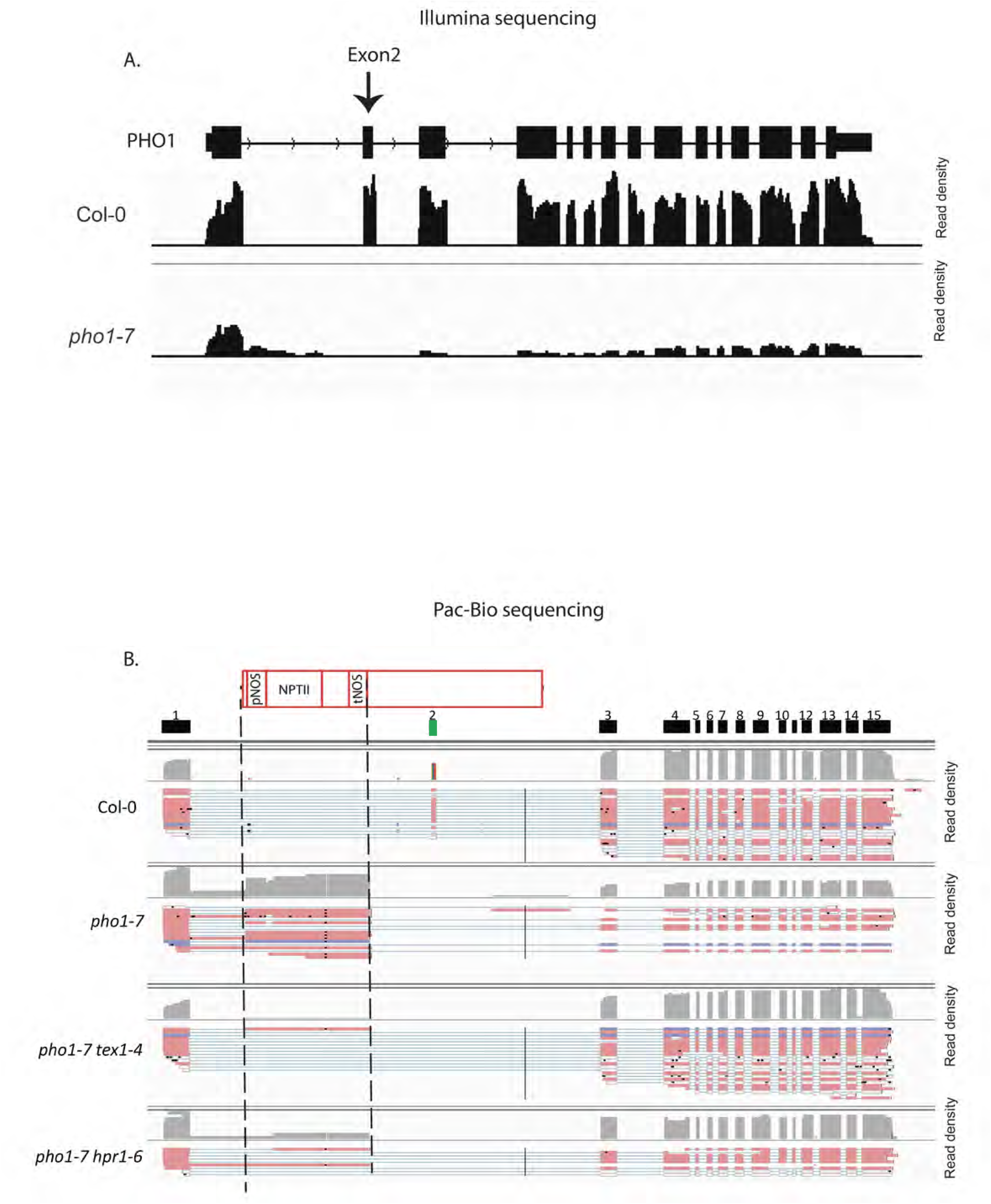
Mapping of *pho1-7* RNA onto the *PHO1* locus. (A) Illumina RNA sequencing reads density graph mapped against the wild-type *PHO1* locus. Note that exon 2 of *PHO1* is missing in the *pho1-7* mutant because of T-DNA insertion while it is present in Col-0. (B) Pac-Bio read density graph showing full length mRNA structure at the *PHO1* locus. The top red box shows the location of the T-DNA in *pho1-7*. pNOS, NOS promoter; NPTII, neomycin phosphotransferase gene; tNOS, NOS terminator. The black boxes below shows the positions of the 15 *PHO1* exons, except for exon 2 which is shown in green. Exon 2 is present in Col-0 but deleted in the *pho1-7*, *pho1-7 tex1-4* and *pho1-7 hpr1-6* mutants as a result of T-DNA insertion. For each genotype, the sequence of independent cDNAs are shown by individual lines (red and blue lines) with the grey areas representing the sequence density in each region.

**S4 Fig.**
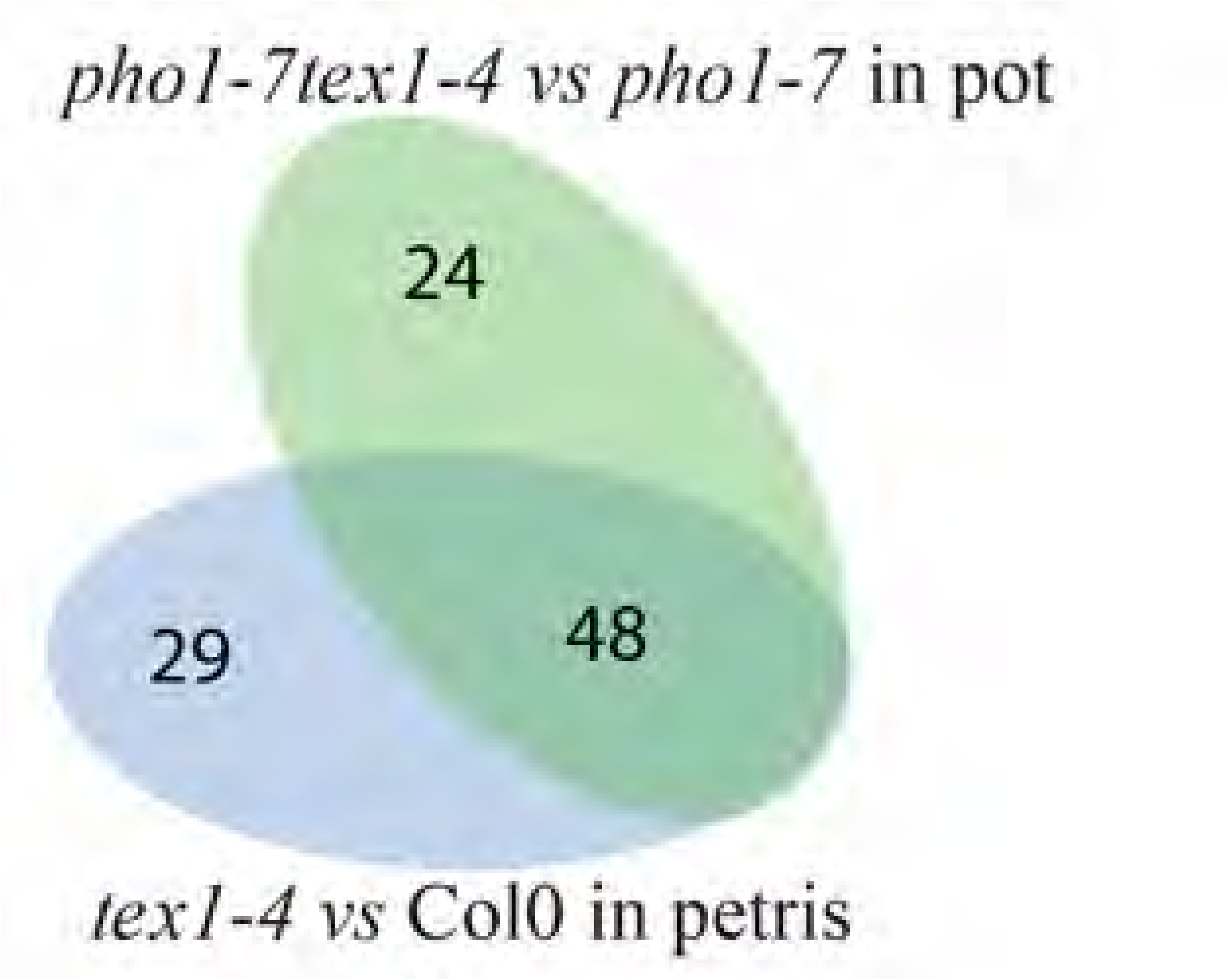
Overlap in genes showing 3’UTR extensions in *tex1-4* mutant grown under various conditions. RNA extracted from roots of *pho1-7 tex1-4* and *pho1-7* plants grown in pots for 4 weeks (green) or *tex1-4* and Col-0 plants grown in petri dishes for 7 days (blue) were used.

**S5 Fig.**
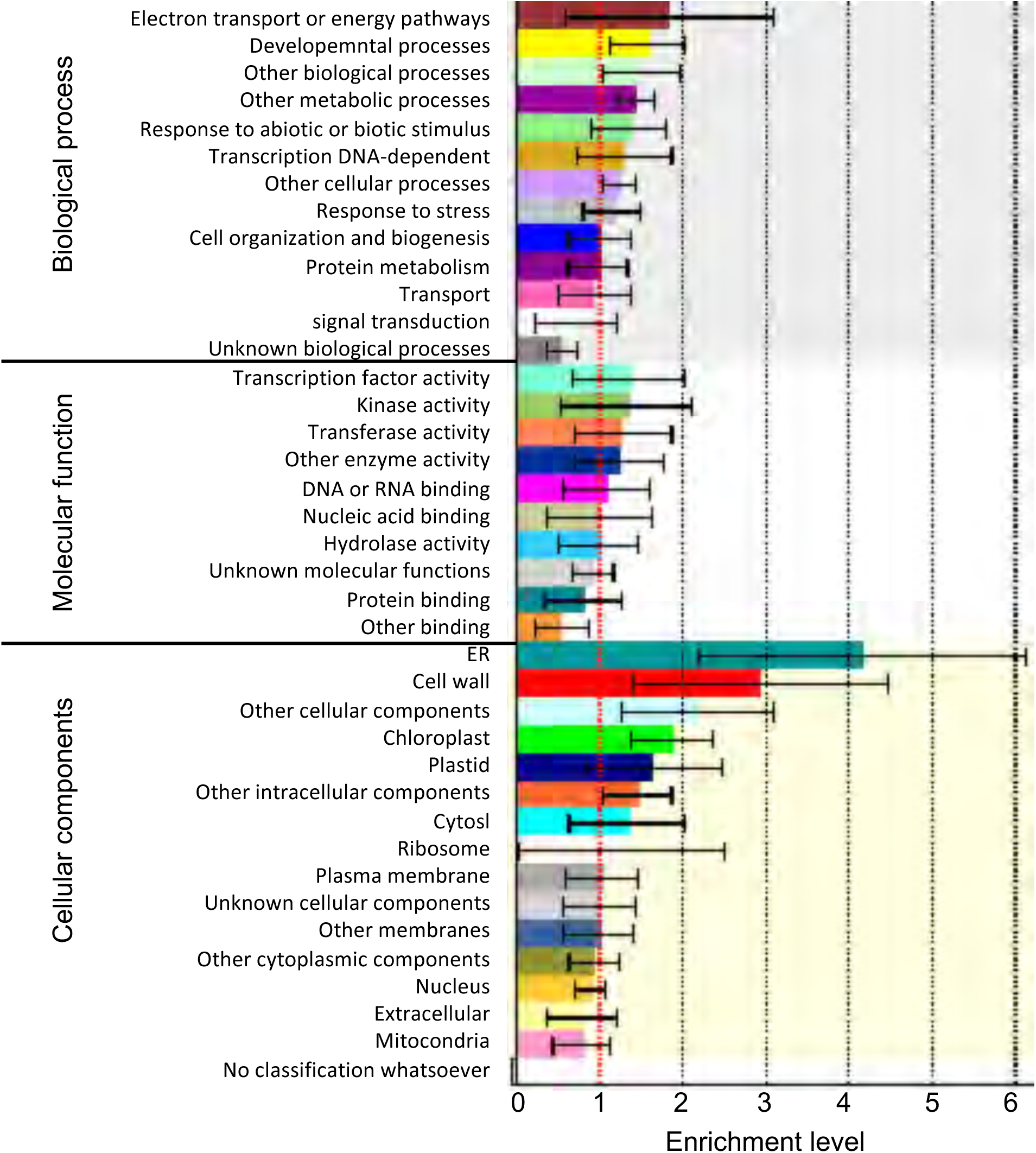
Gene ontology of genes showing 3’UTR extensions in the *hpr1-6* and *tex1-4* **mutants.** The histograms show the fold enrichment of a given gene ontology term.

**Supporting Text S1.**
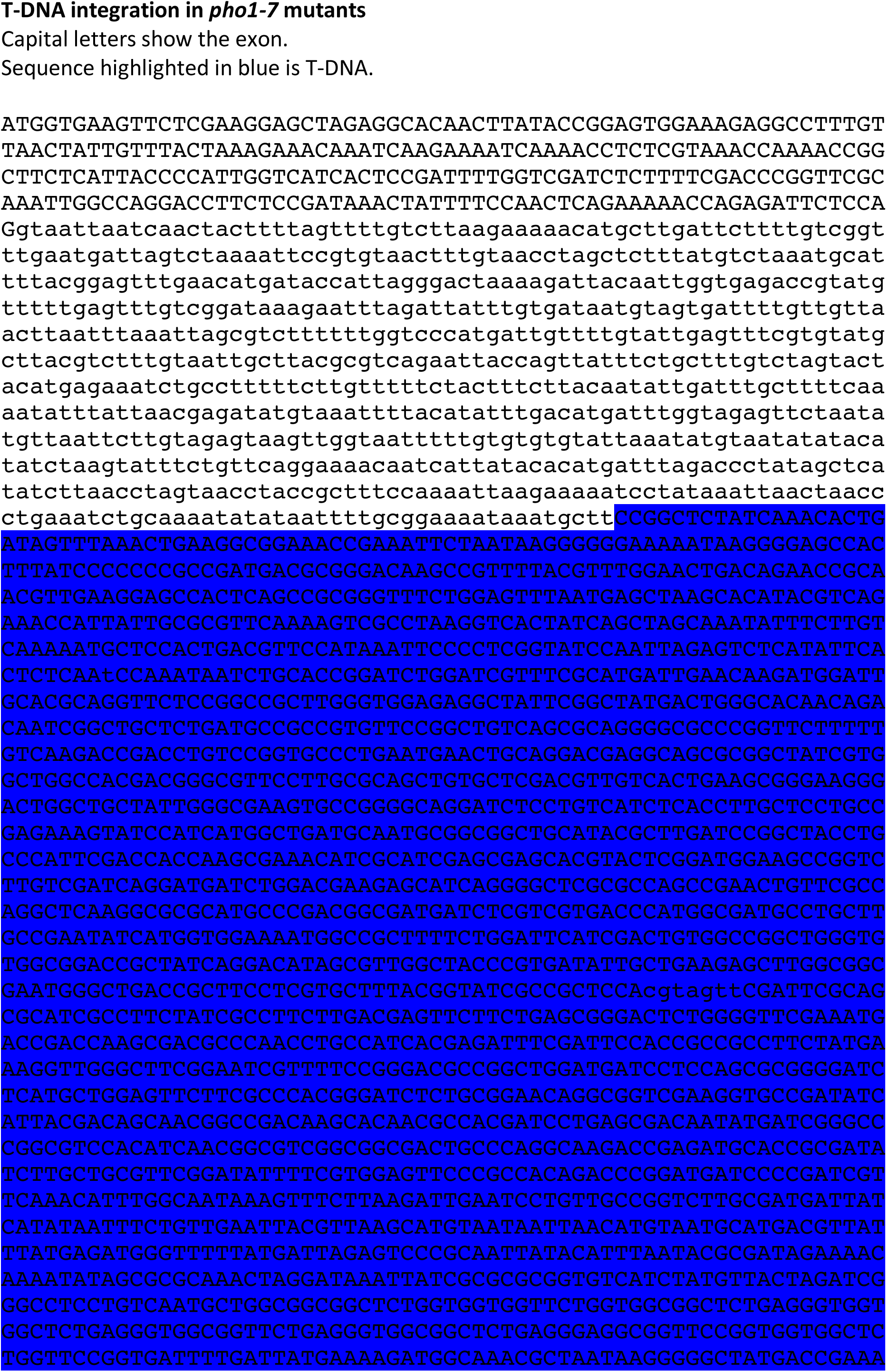
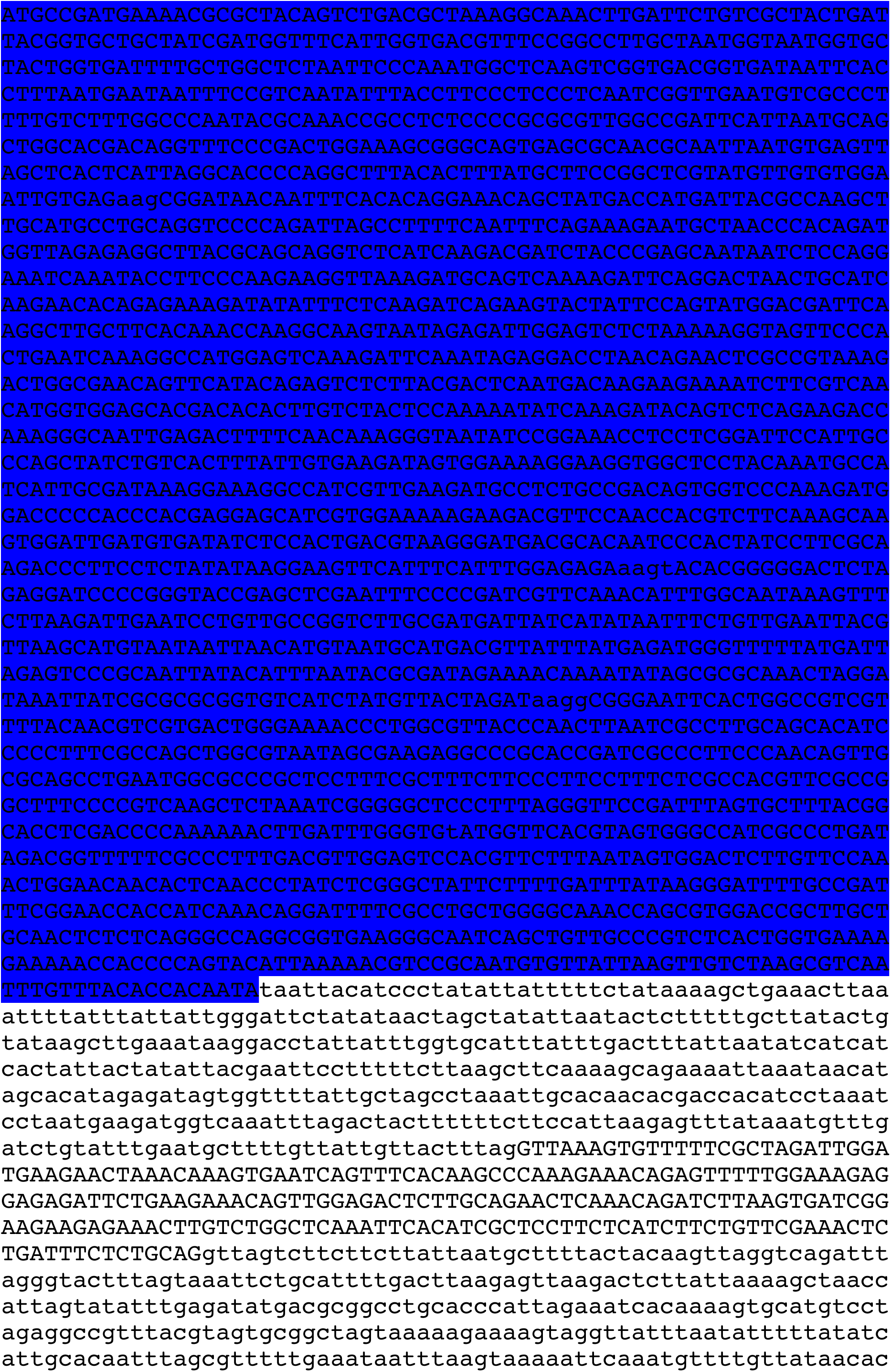
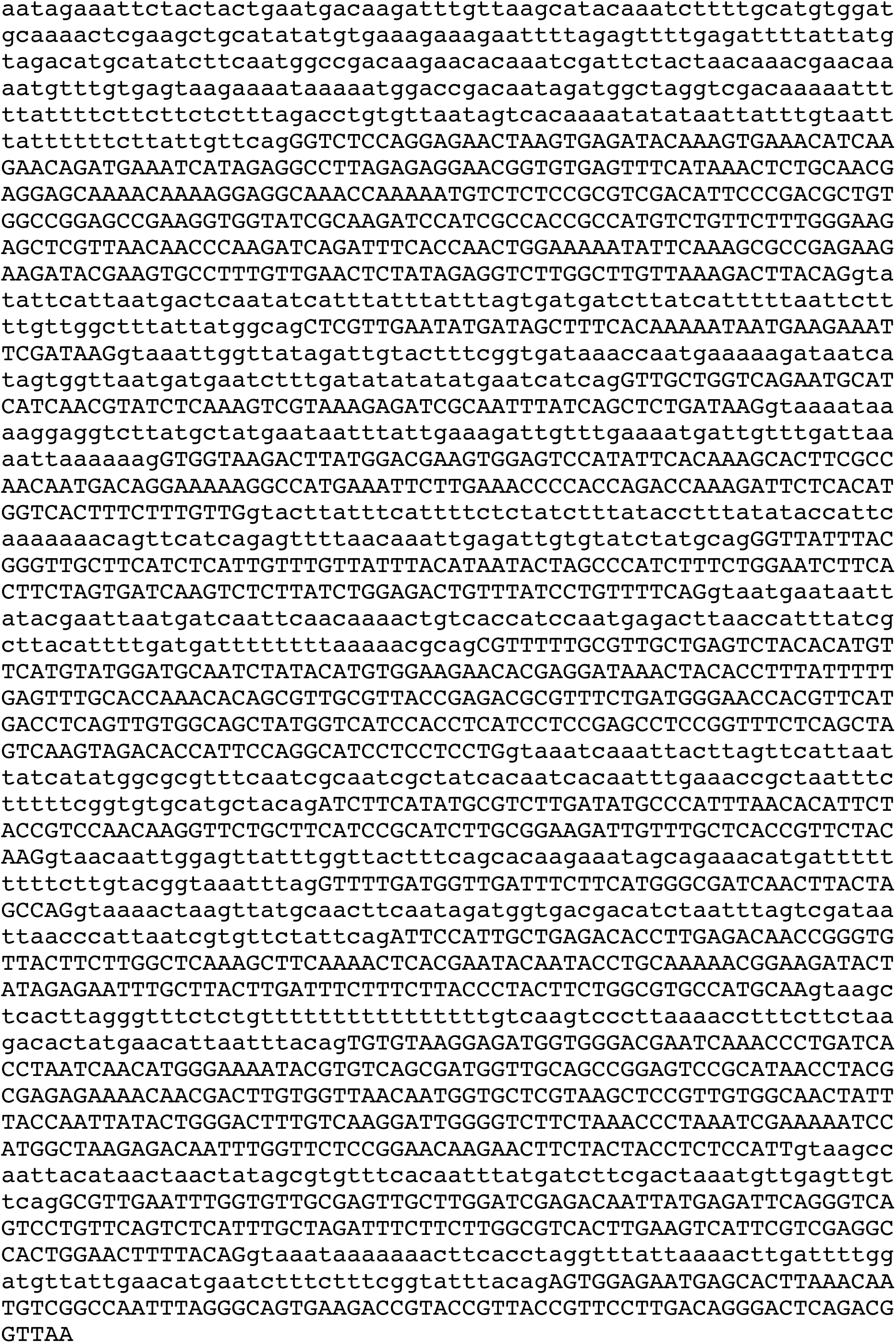
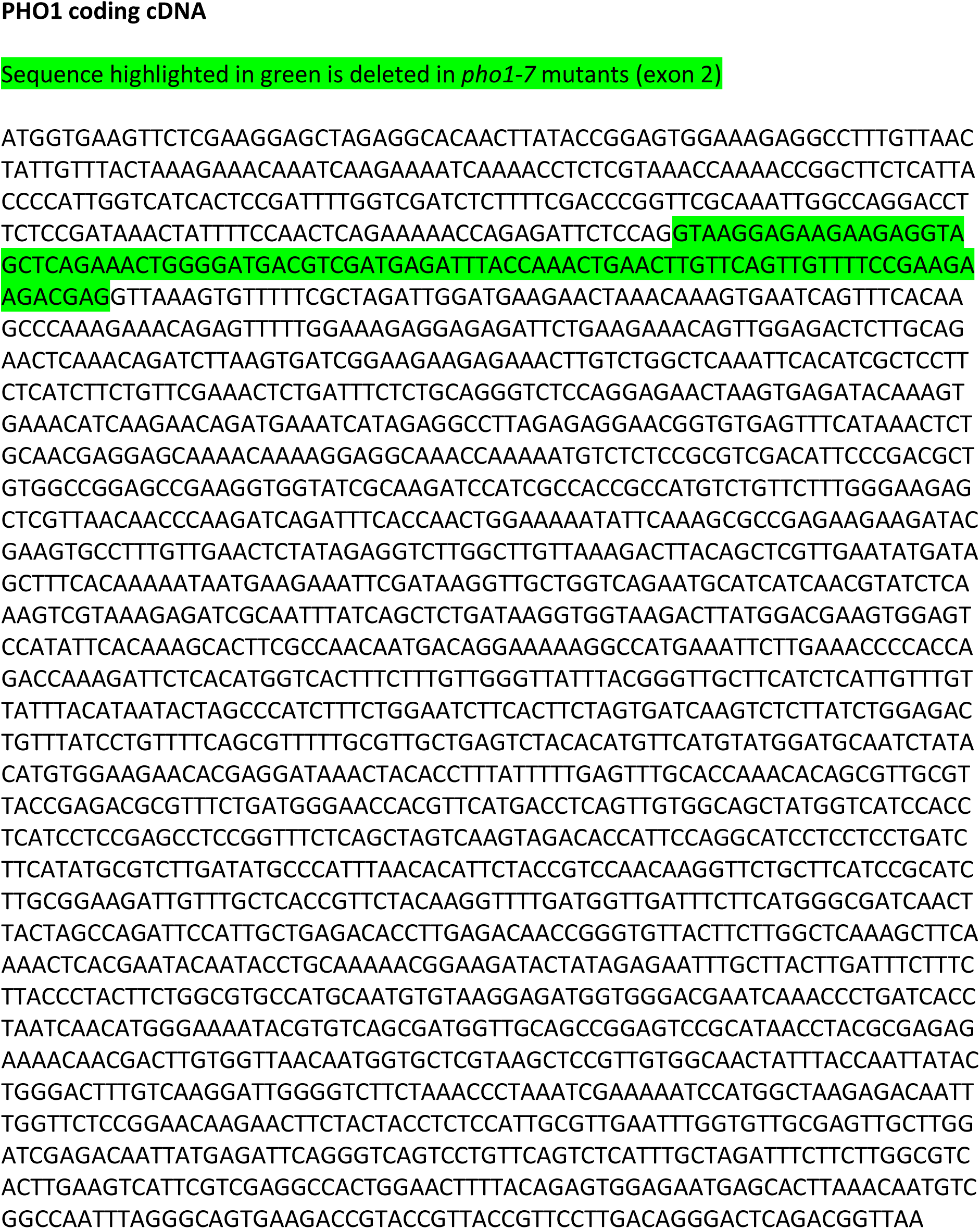
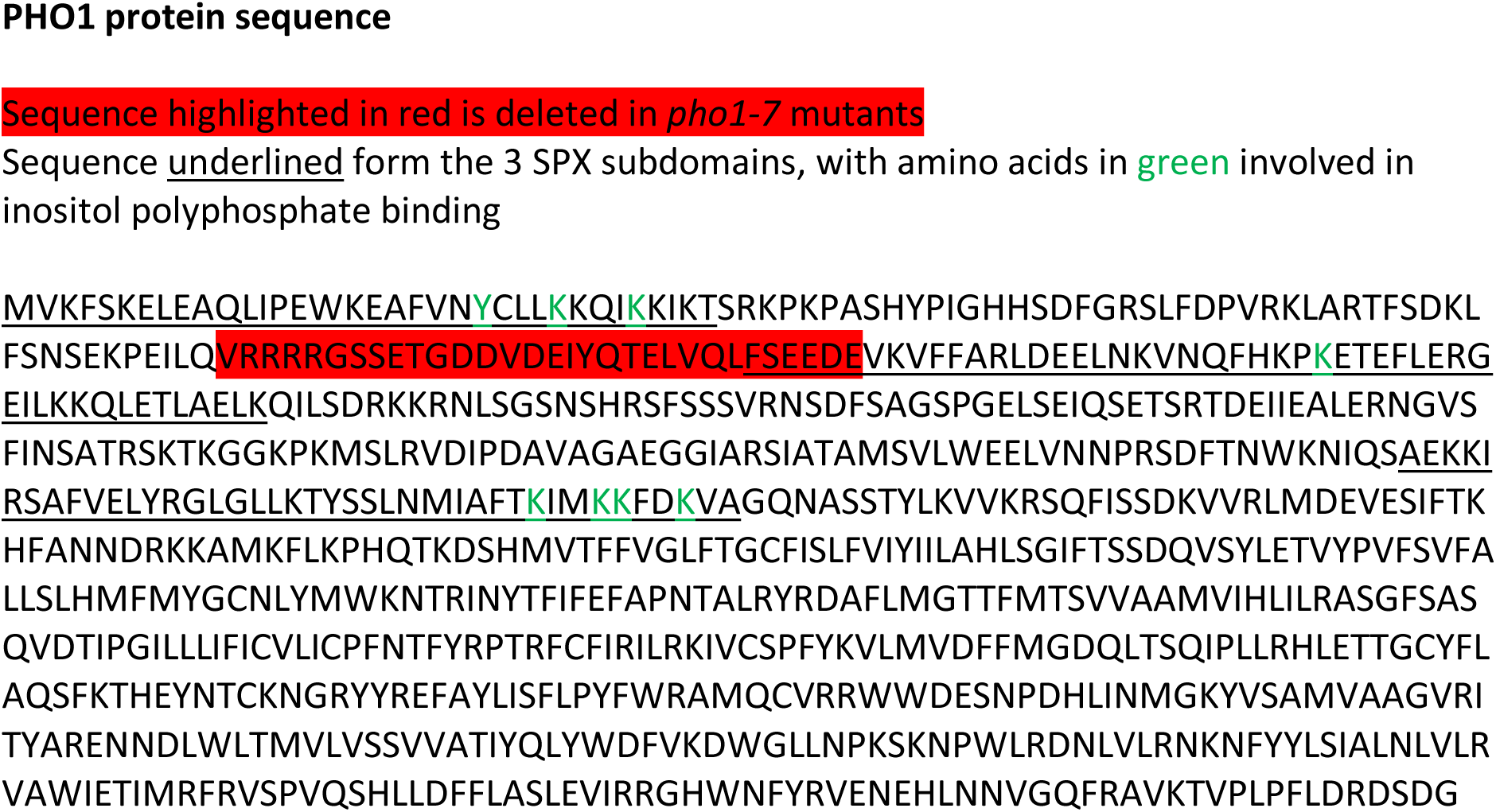
Sequence of the *pho1-7* locus with the production of the truncated PHO1 protein.

**Supporting Table S1.**
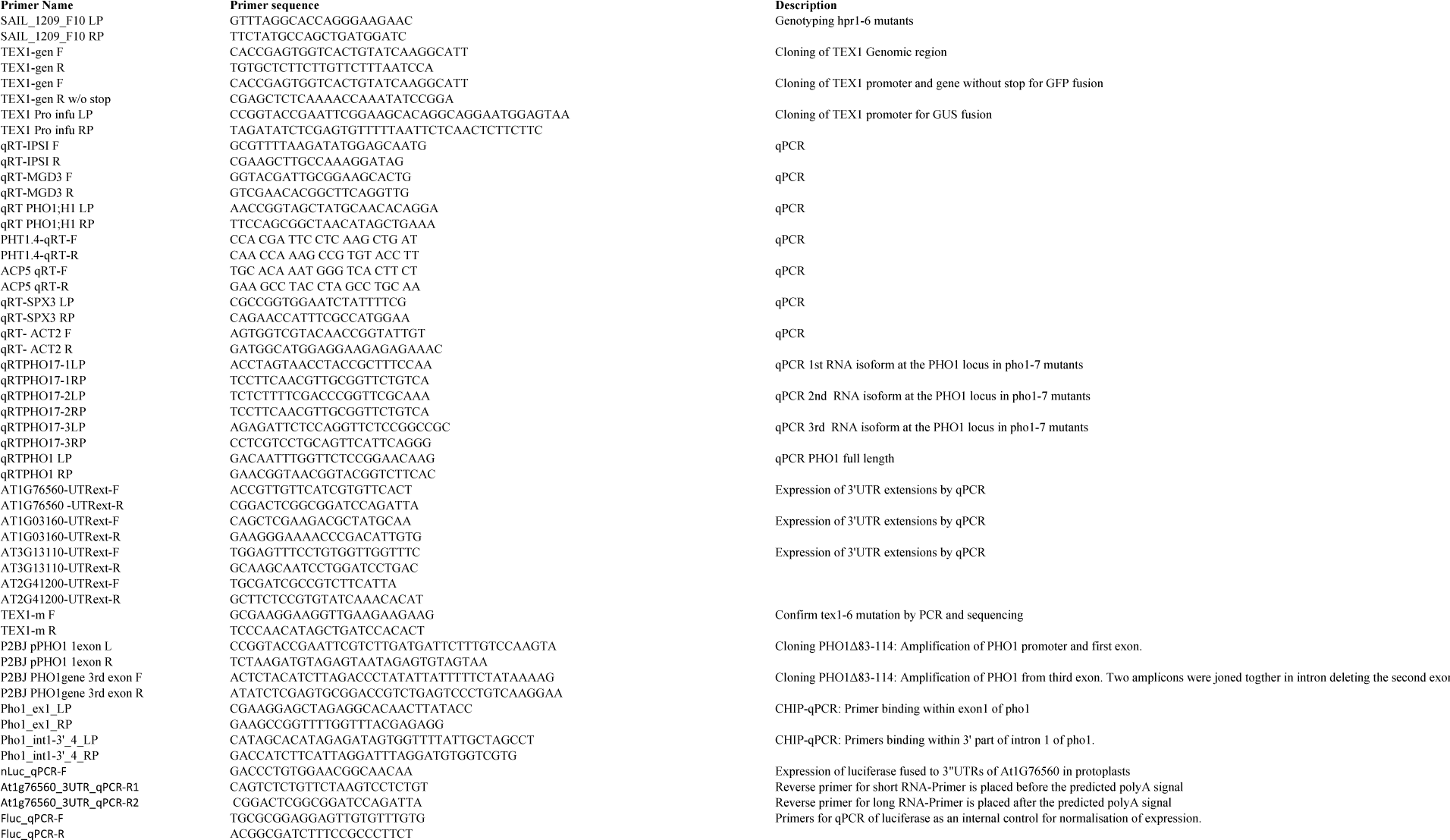
List of oligonucleotides used in this study. Primers Table

## References

1. Chen FX, Smith ER, Shilatifard A. Born to run: control of transcription elongation by RNA polymerase II. Nat Rev Mol Cell Biol. 2018;19(7):464–478.

2. Herzel L, Ottoz DSM, Alpert T, Neugebauer KM. Splicing and transcription touch base: co-transcriptional spliceosome assembly and function. Nat Rev Mol Cell Biol. 2017;18(10):637–650.

3. Richard P, Manley JL. Transcription termination by nuclear RNA polymerases. Genes Dev. 2009;23(11):1247–1269.

4. Hsin J-P, Manley JL. The RNA polymerase II CTD coordinates transcription and RNA processing. Genes Dev. 2012;26(19):2119–2137.

5. Nagarajan VK, Jones CI, Newbury SF, Green PJ. XRN 5’ -> 3’ exoribonucleases: Structure, mechanisms and functions. Biochim Biophys Acta. 2013;1829(6-7):590–603.

6. Chen W, Jia Q, Song Y, Fu H, Wei G, Ni T. Alternative Polyadenylation: Methods, Findings, and Impacts. Genom Proteom Bioinf. 2017;15(5):287–300.

7. Mapendano CK, Lykke-Andersen S, Kjems J, Bertrand E, Jensen TH. Crosstalk between mRNA 3’ End Processing and Transcription Initiation. Mol Cell. 2010;40(3):410–422.

8. Sun H-X, Li Y, Niu Q-W, Chua N-H. Dehydration stress extends mRNA 3’ untranslated regions with noncoding RNA functions in Arabidopsis. Genome Res. 2017;27(8):1427–1436.

9. Katahira J. mRNA export and the TREX complex. Biochim Biophys Acta. 2012;1819(6):507–513.

10. Heath CG, Viphakone N, Wilson SA. The role of TREX in gene expression and disease. Biochem J. 2016;473:2911–2935.

11. Meinel DM, Burkert-Kautzsch C, Kieser A, O’Duibhir E, Siebert M, Mayer A, Cramer P, Soding J, Holstege FCP, Straesser K. Recruitment of TREX to the Transcription Machinery by Its Direct Binding to the Phospho-CTD of RNA Polymerase II. PLoS Genet. 2013;9(11).

12. Gomez-Gonzalez B, Garcia-Rubio M, Bermejo R, Gaillard H, Shirahige K, Marin A, Foiani M, Aguilera A. Genome-wide function of THO/TREX in active genes prevents R-loop-dependent replication obstacles. EMBO J. 2011;30(15):3106–3119.

13. Ehrnsberger HF, Grasser M, Grasser KD. Nucleocytosolic mRNA transport in plants: export factors and their influence on growth and development. J Exp Bot. 2019;doi:10.1093/jxb/erz173.

14. Yelina NE, Smith LM, Jones AME, Patel K, Kelly KA, Baulcombe DC. Putative Arabidopsis THO/TREX mRNA export complex is involved in transgene and endogenous siRNA biosynthesis. Proc Natl Acad Sci USA. 2010;107(31):13948–13953.

15. Jauvion V, Elmayan T, Vaucheret H. The Conserved RNA Trafficking Proteins HPR1 and TEX1 Are Involved in the Production of Endogenous and Exogenous Small Interfering RNA in Arabidopsis. Plant Cell. 2010;22(8):2697–2709.

16. Tao S, Zhang Y, Wang X, Xu L, Fang X, Lu ZJ, Liu D. The THO/TREX Complex Active in miRNA Biogenesis Negatively Regulates Root-Associated Acid Phosphatase Activity Induced by Phosphate Starvation. Plant Physiol. 2016;171(4):2841–2853.

17. Francisco-Mangilet AG, Karlsson P, Kim M-H, Eo HJ, Oh SA, Kim JH, Kulcheski FR, Park SK, Andres Manavella P. THO2, a core member of the THO/TREX complex, is required for microRNA production in Arabidopsis. Plant J. 2015;82(6):1018–1029.

18. Furumizu C, Tsukaya H, Komeda Y. Characterization of EMU, the Arabidopsis homolog of the yeast THO complex member HPR1. RNA. 2010;16(9):1809–1817.

19. Sorensen BB, Ehrnsberger HF, Esposito S, Pfab A, Bruckmann A, Hauptmann J, Meister G, Merkl R, Schubert T, Laengst G et al. The Arabidopsis THO/TREX component TEX1 functionally interacts with MOS11 and modulates mRNA export and alternative splicing events. Plant Mol Biol. 2017;93(3):283–298.

20. Xu C, Zhou X, Wen C-K. HYPER RECOMBINATION1 of the THO/TREX Complex Plays a Role in Controlling Transcription of the REVERSION-TO-ETHYLENE SENSITIVITY1 Gene in Arabidopsis. PLoS Genet. 2015;11(2).

21. Arpat AB, Magliano P, Wege S, Rouached H, Stefanovic A, Poirier Y. Functional expression of PHO1 to the Golgi and *trans*-Golgi network and its role in export of inorganic phosphate. Plant J. 2012;71:479–491.

22. Hamburger D, Rezzonico E, MacDonald-Comber Petétot J, Somerville C, Poirier Y. Identification and characterization of the *Arabidopsis PHO1* gene involved in phosphate loading to the xylem. Plant Cell. 2002;14:889–902.

23. Poirier Y, Thoma S, Somerville C, Schiefelbein J. A mutant of *Arabidopsis* deficient in xylem loading of phosphate. Plant Physiol. 1991;97:1087–1093.

24. Stefanovic A, Arpat AB, Bligny R, Gout E, Vidoudez C, Bensimon M, Poirier Y. Overexpression of PHO1 in Arabidopsis leaves reveals its role in mediating phosphate efflux. Plant J. 2011;66:689–699.

25. Rouached H, Stefanovic A, Secco D, Arpat AB, Gout E, Bligny R, Poirier Y. Uncoupling phosphate deficiency from its major effects on growth and transcriptome via PHO1 expression in Arabidopsis. Plant J. 2011;65:557–570.

26. Secco D, Baumann A, Poirier Y. Charcaterization of the rice *PHO1* gene family reveals a key role for *OsPHO1;2* in phosphate homeostasis and the evolution of a distinct clade in dicotyledons. Plant Physiol. 2010;152:1693–1704.

27. Wege S, Khan GA, Jung J-Y, Vogiatzaki E, Pradervand S, Aller I, Meyer AJ, Poirier Y. The EXS domain of PHO1 participates in the response of shoots to phosphate deficiency via a root-to-shoot signal. Plant Physiol. 2016;170(1):385–400.

28. Peragine A, Yoshikawa M, Wu G, Albrecht HL, Poethig RS. SGS3 and SGS2/SDE1/RDR6 are required for juvenile development and the production of trans-acting siRNAs in Arabidopsis. Genes Dev. 2004;18(19):2368–2379.

29. Ronemus M, Vaughn MW, Martienssen RA. MicroRNA-targeted and small interfering RNA-mediated mRNA degradation is regulated by Argonaute, Dicer, and RNA-dependent RNA polymerase in Arabidopsis. Plant Cell. 2006;18(7):1559–1574.

30. Loke JC, Stahlberg EA, Strenski DG, Haas BJ, Wood PC, Li QQ. Compilation of mRNA polyadenylation signals in Arabidopsis revealed a new signal element and potential secondary structures. Plant Physiol. 2005;138(3):1457–1468.

31. Sherstnev A, Duc C, Cole C, Zacharaki V, Hornyik C, Ozsolak F, Milos PM, Barton GJ, Simpson GG. Direct sequencing of Arabidopsis thaliana RNA reveals patterns of cleavage and polyadenylation. Nat Struct Mol Biol. 2012;19(8):845–852.

32. Wu XH, Liu M, Downie B, Liang C, Ji GL, Li QQ, Hunt AG. Genome-wide landscape of polyadenylation in Arabidopsis provides evidence for extensive alternative polyadenylation. Proc Natl Acad Sci USA. 2011;108(30):12533–12538.

33. Doan Duy Hai T, Saran S, Williamson AJK, Pierce A, Dittrich-Breiholz O, Wiehlmann L, Koch A, Whetton AD, Tamura T. THOC5 controls 3’ end-processing of immediate early genes via interaction with polyadenylation specific factor 100 (CPSF100). Nucleic Acids Res. 2014;42(19):12249–12260.

34. Katahira J, Okuzaki D, Inoue H, Yoneda Y, Maehara K, Ohkawa Y. Human TREX component Thoc5 affects alternative polyadenylation site choice by recruiting mammalian cleavage factor I. Nucleic Acids Res. 2013;41(14):7060–7072.

35. Pak V, Eifler TT, Jaeger S, Krogan NJ, Fujinaga K, Peterlin BM. CDK11 in TREX/THOC Regulates HIV mRNA 3’ End Processing. Cell Host Microbe. 2015;18(5):560–570.

36. Pan H, Liu S, Tang D. HPR1, a component of the THO/TREX complex, plays an important role in disease resistance and senescence in Arabidopsis. Plant J. 2012;69(5):831–843.

37. Doll S, Kuhlmann M, Rutten T, Mette MF, Scharfenberg S, Petridis A, Berreth DC, Mock HP. Accumulation of the coumarin scopolin under abiotic stress conditions is mediated by the Arabidopsis thaliana THO/TREX complex. Plant J. 2018;93(3):431–444.

38. Su Z, Zhao L, Zhao Y, Li S, Won S, Cai H, Wang L, Li Z, Chen P, Qin Y et al. The THO Complex Non-Cell-Autonomously Represses Female Germline Specification through the TAS3-ARF3 Module. Curr Biol. 2017;27(11):1597-+.

39. Tian B, Hu J, Zhang HB, Lutz CS. A large-scale analysis of mRNA polyadenylation of human and mouse genes. Nucleic Acids Res. 2005;33(1):201–212.

40. Depicker A, Stachel S, Dhaese P, Zambryski PC, Goodman H. Nopaline synthase: transcript mapping and DNA sequence. J Mol Appl Genet. 1982;6:561–573.

41. Beyene G, Buenrostro-Nava MT, Damaj MB, Gao SJ, Molina J, Mirkov TE. Unprecedented enhancement of transient gene expression from minimal cassettes using a double terminator. Plant Cell Rep. 2011;30(1):13–25.

42. Nagaya S, Kawamura K, Shinmyo A, Kato K. The HSP Terminator of Arabidopsis thaliana Increases Gene Expression in Plant Cells. Plant Cell Physiol. 2010;51(2):328–332.

43. Richter LJ, Thanavala Y, Arntzen CJ, Mason HS. Production of hepatitis B surface antigen in transgenic plants for oral immunization. Nat Biotechnol. 2000;18(11):1167–1171.

44. Rosenthal SH, Diamos AG, Mason HS. An intronless form of the tobacco extensin gene terminator strongly enhances transient gene expression in plant leaves. Plant Mol Biol. 2018;96(4-5):429–443.

45. Mayr C. Regulation by 3 ‘-untranslated regions. Ann Rev Genet. 2017;51:171–194.

46. Kindgren P, Ard R, Ivanov M, Marquardt S. Transcriptional read-through of the long non-coding RNA SVALKA governs plant cold acclimation. Nature Comm. 2018;9.

47. Cuerda-Gil D, Slotkin RK. Non-canonical RNA-directed DNA methylation. Nature Plants. 2016;2(11):e16163.

48. Mathieu O, Bouche N. Interplay between chromatin and RNA processing. Curr Opin Plant Biol. 2014;18:60–65.

49. Jung J-Y, Ried MK, Hothorn M, Poirier Y. Control of plant phosphate homeostasis by inositol pyrophosphates and the SPX domain. Curr Opin Biotechnol. 2018;49:156–162.

50. Wild R, Gerasimaite R, Jung J-Y, Truffault V, Pavlovic I, Schmidt A, Saiardi A, Jessen HJ, Poirier Y, Hothorn M et al. Control of eukaryotic phosphate homeostasis by inositol polyphosphate sensor domains. Science. 2016;352(6288):986–990.

51. Liu T-Y, Huang T-K, Tseng C-Y, Lai Y-S, Lin S-I, Lin W-Y, Chen J-W, Chiou T-J. PHO2-dependent degradation of PHO1 modulates phosphate homeostasis in Arabidopsis. Plant Cell. 2012;24(5):2168–2183.

52. Khan GA, Vogiatzaki E, Glauser G, Poirier Y. Phosphate Deficiency Induces the Jasmonate Pathway and Enhances Resistance to Insect Herbivory. Plant Physiol. 2016;171(1):632–644.

53. Ames BN. Assay of inorganic phosphate, total phosphate and phosphatases. Methods Enzymol. 1966;8:115–118.

54. Curtis MD, Grossniklaus U. A gateway cloning vectors set for high-throughput functional analysis of genes in planta. Plant Physiol. 2003;133:462–469.

55. Clough SJ, Bent AF. Floral dip: a simplified method for *Agrobacterium*-mediated transformation of *Arabidopsis thaliana*. Plant J. 1998;6:735–743.

56. Deforges J, Reis RS, Jacquet P, Sheppard S, Geadekar VP, Hart-Smith G, Tanzer A, Hofacker IL, Iseli C, Xenarios I et al. Control of cognate mRNA translation by *cis*-natural antisense. Plant Physiol. 2019;180(1):305–322.

57. Kim D, Landmead B, Salzberg SL. HISAT: a fast spliced aligner with low memory requirements. Nat Methods. 2015;12(4):357–360.

58. Anders S, Pyl PT, Huber W. HTSeq-a Python framework to work with high-throughput sequencing data. Bioinformatics. 2015;31(2):166–169.

59. Love M, Huber W, Anders S. Moderated estimation of fold change and dispersion for RNA-seq data with DESeq2. Genome Biol. 2014;15:550.

60. Robinson JT, Thorvaldsdottir H, Winckler W, Guttman M, Lander ES, Getz G, Mesirov JP. Integrative genomics viewer. Nat Biotechnol. 2011;29(1):24–26.

61. Yoo S-D, Cho Y-H, Sheen J. Arabidopsis mesophyll protoplasts: a versatile cell system for transient gene expression analysis. Nat Protoc. 2007;2(7):1565–1572.

