## Supplementary figures and images for "The transcription and export complex THO/TREX contributes to transcription termination in plants"

### S1 Figure

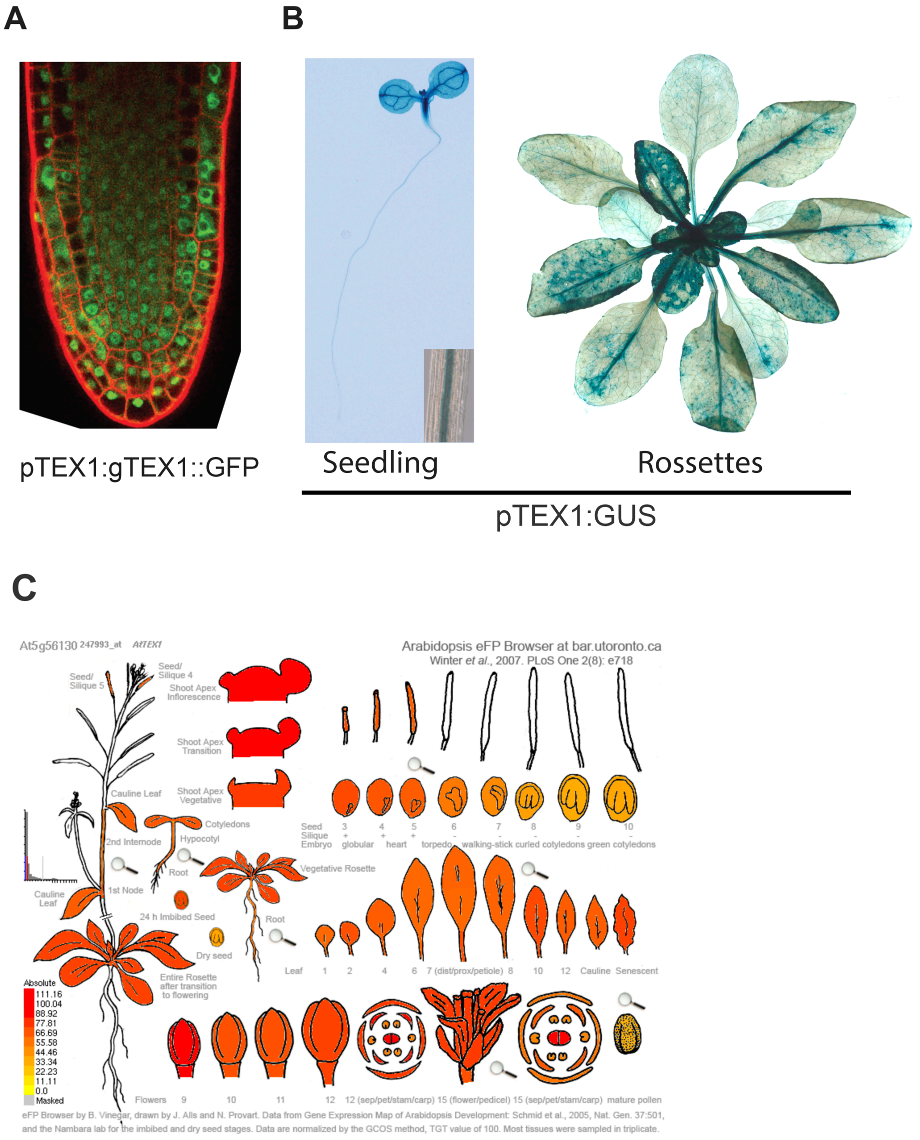

### S2 Figure

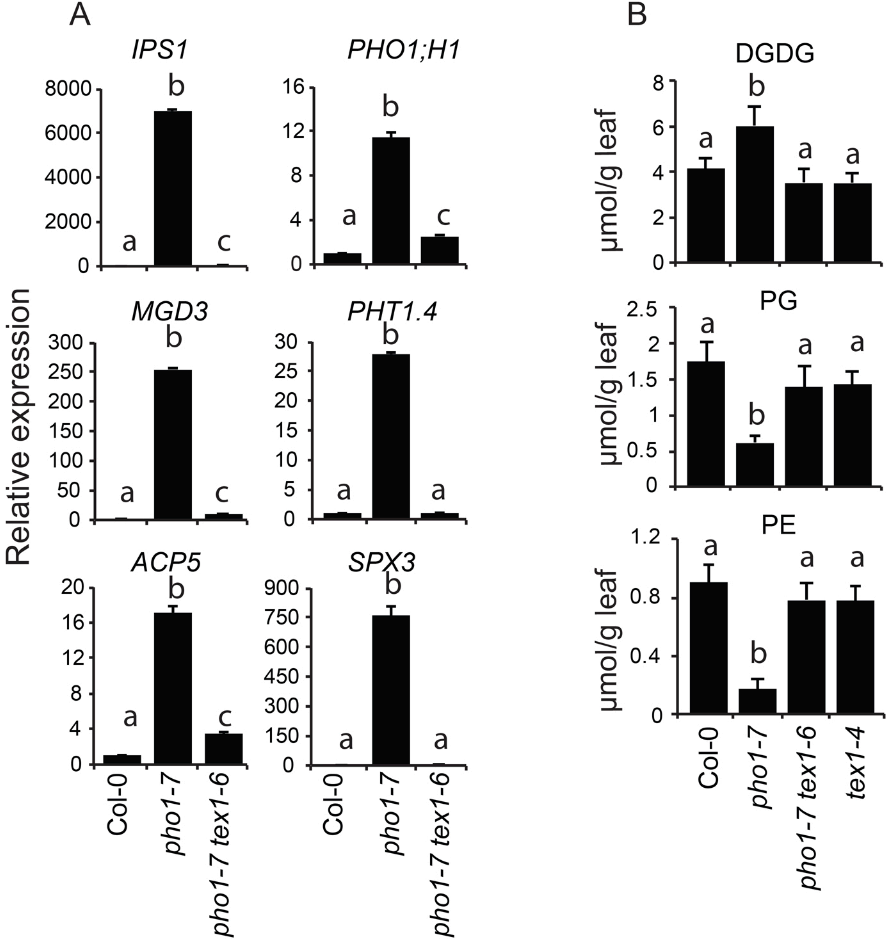

### S3 Figure

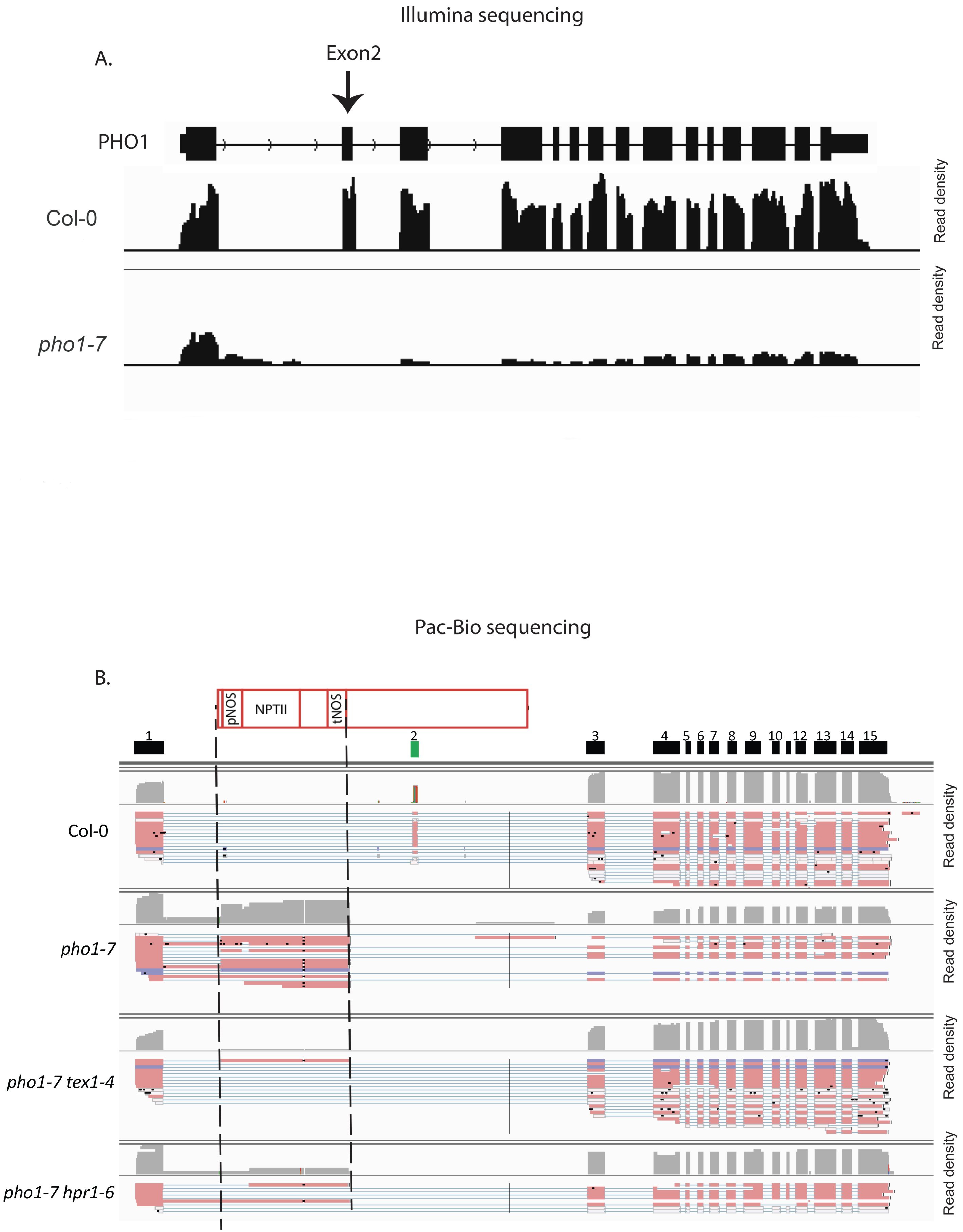

### S3 Figure

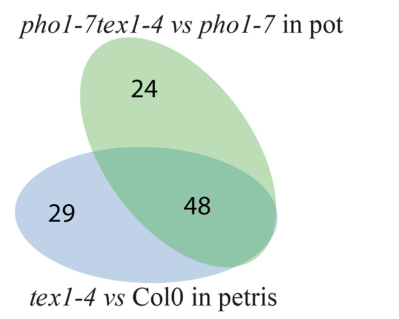

### S5 Figure

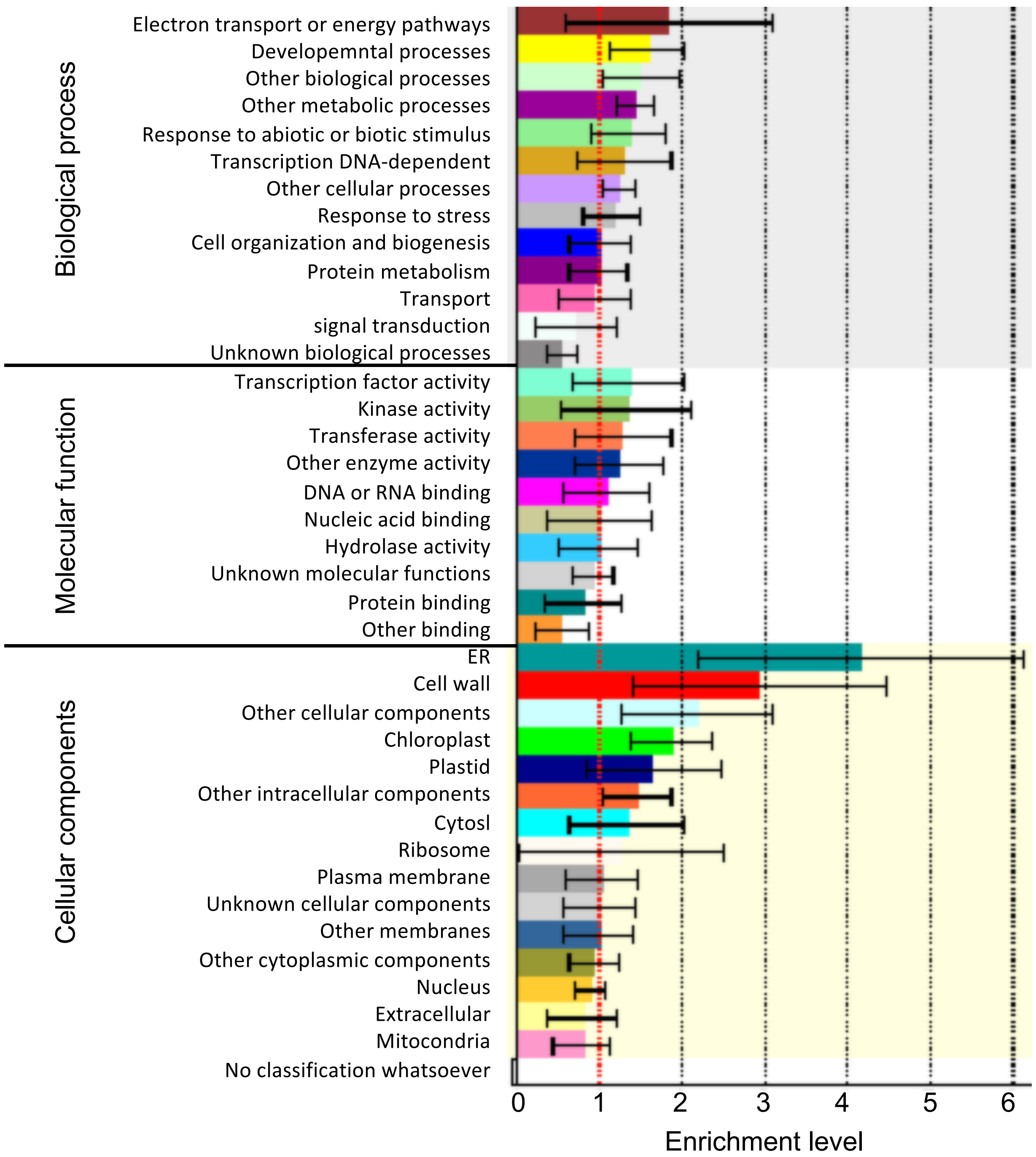
