## Supplementary material for "The transcription and export complex THO/TREX contributes to transcription termination in plants": Table

| Primer Name | Primer sequence | Description |
|---|---|---|
| SAIL_1209_F10 LP | GTTTAGGCACCAGGGAAGAAC | Genotyping hpr1-6 mutants |
| SAIL_1209_F10 RP | TTCTATGCCAGCTGATGGATC | |
| TEX1-gen F | CACCGAGTGGTCACTGTATCAAGGCATT | Cloning of TEX1 Genomic region |
| TEX1-gen R | TGTGCTCTTCTTGTTCTTTAATCCA | |
| TEX1-gen F | CACCGAGTGGTCACTGTATCAAGGCATT | Cloning of TEX1 promoter and gene without stop for GFP fusion |
| TEX1-gen R w/o stop | CGAGCTCTCAAAACCAAATATCCGGA | |
| TEX1 Pro infu LP | CCGGTACCGAATTCGGAAGCACAGGCAGGAATGGAGTAA | Cloning of TEX1 promoter for GUS fusion |
| TEX1 Pro infu RP | TAGATATCTCGAGTGTTTTTAATTCTCAACTCTTCTTC | |
| qRT-IPSI F | GCGTTTTAAGATATGGAGCAATG | qPCR |
| qRT-IPSI R | CGAAGCTTGCCAAAGGATAG | |
| qRT-MGD3 F | GGTACGATTGCGGAAGCACTG | qPCR |
| qRT-MGD3 R | GTCGAACACGGCTTCAGGTTG | |
| qRT PHO1;H1 LP | AACCGGTAGCTATGCAACACAGGA | qPCR |
| qRT PHO1;H1 RP | TTCCAGCGGCTAACATAGCTGAAA | |
| PHT1.4-qRT-F | CCA CGA TTC CTC AAG CTG AT | qPCR |
| PHT1.4-qRT-R | CAA CCA AAG CCG TGT ACC TT | |
| ACP5 qRT-F | TGC ACA AAT GGG TCA CTT CT | qPCR |
| ACP5 qRT-R | GAA GCC TAC CTA GCC TGC AA | |
| qRT-SPX3 LP | CGCCGGTGGAATCTATTTTCG | qPCR |
| qRT-SPX3 RP | CAGAACCATTTCGCCATGGAA | |
| qRT- ACT2 F | AGTGGTCGTACAACCGGTATTGT | qPCR |
| qRT- ACT2 R | GATGGCATGGAGGAAGAGAGAAAC | |
| qRTPHO17-1LP | ACCTAGTAACCTACCGCTTTCCAA | qPCR 1st RNA isoform at the PHO1 locus in pho1-7 mutants |
| qRTPHO17-1RP | TCCTTCAACGTTGCGGTTCTGTCA | |
| qRTPHO17-2LP | TCTCTTTTCGACCCGGTTCGCAAA | qPCR 2nd RNA isoform at the PHO1 locus in pho1-7 mutants |
| qRTPHO17-2RP | TCCTTCAACGTTGCGGTTCTGTCA | |
| qRTPHO17-3LP | AGAGATTCTCCAGGTTCTCCGGCCGC | qPCR 3rd RNA isoform at the PHO1 locus in pho1-7 mutants |
| qRTPHO17-3RP | CCTCGTCCTGCAGTTCATTCAGGG | |
| qRTPHO1 LP | GACAATTTGGTTCTCCGGAACAAG | qPCR PHO1 full length |
| qRTPHO1 RP | GAACGGTAACGGTACGGTCTTCAC | |
| AT1G76560-UTRext-F | ACCGTTGTTCATCGTGTTCACT | Expression of 3'UTR extensions by qPCR |
| AT1G76560 -UTRext-R | CGGACTCGGCGGATCCAGATTA | |
| AT1G03160-UTRext-F | CAGCTCGAAGACGCTATGCAA | Expression of 3'UTR extensions by qPCR |
| AT1G03160-UTRext-R | GAAGGGAAAACCCGACATTGTG | |
| AT3G13110-UTRext-F | TGGAGTTTCCTGTGGTTGGTTTC | Expression of 3'UTR extensions by qPCR |
| AT3G13110-UTRext-R | GCAAGCAATCCTGGATCCTGAC | |
| AT2G41200-UTRext-F | TGCGATCGCCGTCTTCATTA | |
| AT2G41200-UTRext-R | GCTTCTCCGTGTATCAAACACAT | |
| TEX1-m F | GCGAAGGAAGGTTGAAGAAGAAG | Confirm tex1-6 mutation by PCR and sequencing |
| TEX1-m R | TCCCAACATAGCTGATCCACACT | |
| P2BJ pPHO1 1exon L | CCGGTACCGAATTCGTCTTGATGATTCTTTGTCCAAGTA | Cloning PHO1Δ83-114: Amplification of PHO1 promoter and first exon. |
| P2BJ pPHO1 1exon R | TCTAAGATGTAGAGTAATAGAGTGTAGTAA | |
| P2BJ PHO1gene 3rd exon F | ACTCTACATCTTAGACCCTATATTATTTTTCTATAAAAG | Cloning PHO1Δ83-114: Amplification of PHO1 from third exon. Two amplicons were joned togther in intron deleting the second exo[illegible] |
| P2BJ PHO1gene 3rd exon R | ATATCTCGAGTGCGGACCGTCTGAGTCCCTGTCAAGGAA | |
| Pho1_ex1_LP | CGAAGGAGCTAGAGGCACAACTTATACC | CHIP-qPCR: Primer binding within exon1 of pho1 |
| Pho1_ex1_RP | GAAGCCGGTTTTGGTTTACGAGAGG | |
| Pho1_int1-3'_4_LP | CATAGCACATAGAGATAGTGGTTTTATTGCTAGCCT | CHIP-qPCR: Primers binding within 3' part of intron 1 of pho1. |
| Pho1_int1-3'_4_RP | GACCATCTTCATTAGGATTTAGGATGTGGTCGTG | |
| nLuc_qPCR-F | GACCCTGTGGAACGGCAACAA | Expression of luciferase fused to 3"UTRs of At1G76560 in protoplasts |
| At1g76560_3UTR_qPCR-R1 | CAGTCTCTGTTCTAAGTCCTCTGT | Reverse primer for short RNA-Primer is placed before the predicted polyA signal |
| At1g76560_3UTR_qPCR-R2 | CGGACTCGGCGGATCCAGATTA | Reverse primer for long RNA-Primer is placed after the predicted polyA signal |
| Fluc_qPCR-F | TGCGCGGAGGAGTTGTGTTTGTG | Primers for qPCR of luciferase as an internal control for normalisation of expression. |
| Fluc_qPCR-R | ACGGCGATCTTTCCGCCCTTCT | |
