## Supplementary material for "The transcription and export complex THO/TREX contributes to transcription termination in plants": S1 Text

Supplementary Text S1. Sequence of the *pho1-7* locus with the production of the truncated PHO1 protein.

**T-DNA integration in *pho1-7* mutants**

Capital letters show the exon.
Sequence highlighted in blue is T-DNA.

ATGGTGAAGTTCTCGAAGGAGCTAGAGGCACAACTTATACCGGAGTGGAAAGAGGCCTTTGT
TAACTATTGTTTACTAAAGAAACAAATCAAGAAAATCAAAACCTCTCGTAAACCAAAACCGG
CTTCTCATTACCCCATTGGTCATCACTCCGATTTTGGTCGATCTCTTTTCGACCCGGTTCGC
AAATTGGCCAGGACCTTCTCCGATAAACTATTTTCCAACTCAGAAAAACCAGAGATTCTCCA
Ggtaattaatcaactacttttagttttgtcttaagaaaaacatgcttgattcttttgtcggt
ttgaatgattagtctaaaattccgtgtaactttgtaacctagctctttatgtctaaatgcat
tttacggagtttgaacatgataccattagggactaaaagattacaattggtgagaccgtatg
tttttgagtttgtcggataaagaatttagattatttgtgataatgtagtgattttgttgtta
acttaatttaaattagcgtctttttttggtcccatgattgttttgtattgagtttcgtgtatg
cttacgtctttgtaattgcttacgcgtcagaattaccagttatttctgctttgtctagtact
acatgagaaatctgcctttttcttgtttttctactttcttacaatattgatttgcttttcaa
aatatttattaacgagatatgtaaattttacatatttgacatgatttggtagagttctaata
tgttaattcttgtagagtaagttggtaatttttgtgtgtgtattaaatatgtaatatataca
tatctaagtatttctgttcaggaaaacaatcattatacacatgatttagaccctatagctca
tatcttaacctagtaacctaccgctttccaaaattaagaaaaatcctataaattaactaacc
ctgaaatctgcaaaatatataattttgcggaaaataaatgcttCCGGCTCTATCAAACACTG
ATAGTTTAAACTGAAGGCGGAAACCGAAATTCTAATAAGGGGGAAAAATAAGGGGAGCCAC
TTTATCCCCCCCGCCGATGACGCGGGACAAGCCGTTTTACGTTTGGAACTGACAGAACCGCA
ACGTTGAAGGAGCCACTCAGCCGCGGGTTTCTGGAGTTTAATGAGCTAAGCACATACGTCAG
AAACCATTATTGCGCGTTCAAAAGTCGCCTAAGGTCACTATCAGCTAGCAAATATTTCTTGT
CAAAAATGCTCCACTGACGTTCCATAAATTCCCCTCGGTATCCAATTAGAGTCTCATATTCA
CTCTCAAtCCAAATAATCTGCACCGGATCTGGATCGTTTCGCATGATTGAACAAGATGGATT
GCACGCAGGTTCTCCGGCCGCTTGGGTGGAGAGGCTATTCGGCTATGACTGGGCACAACAGA
CAATCGGCTGCTCTGATGCCGCCGTGTTCCGGCTGTCAGCGCAGGGGCGCCCGGTTCTTTTT
GTCAAGACCGACCTGTCCGGTGCCCTGAATGAACTGCAGGACGAGGCAGCGCGGCTATCGTG
GCTGGCCACGACGGGCGTTCCTTGCGCAGCTGTGCTCGACGTTGTCACTGAAGCGGGAAGGG
ACTGGCTGCTATTGGGCGAAGTGCCGGGGCAGGATCTCCTGTCATCTCACCTTGCTCCTGCC
GAGAAAGTATCCATCATGGCTGATGCAATGCGGCGGCTGCATACGCTTGATCCGGCTACCTG
CCCATTCGACCACCAAGCGAAACATCGCATCGAGCGAGCACGTACTCGGATGGAAGCCGGTC
TTGTCGATCAGGATGATCTGGACGAAGAGCATCAGGGGCTCGCGCCAGCCGAACTGTTCGCC
AGGCTCAAGGCGCGCATGCCCGACGGCGATGATCTCGTCGTGACCCATGGCGATGCCTGCTT
GCCGAATATCATGGTGGAAAATGGCCGCTTTTCTGGATTCATCGACTGTGGCCGGCTGGGTG
TGGCGGACCGCTATCAGGACATAGCGTTGGCTACCCGTGATATTGCTGAAGAGCTTGGCGGC
GAATGGGCTGACCGCTTCCTCGTGCTTTACGGTATCGCCGCTCCAcgtagttCGATTCGCAG
CGCATCGCCTTCTATCGCCTTCTTGACGAGTTCTTCTGAGCGGGACTCTGGGGTTCGAAATG
ACCGACCAAGCGACGCCCAACCTGCCATCACGAGATTTCGATTCCACCGCCGCCTTCTATGA
AAGGTTGGGCTTCGGAATCGTTTTCCGGGACGCCGGCTGGATGATCCTCCAGCGCGGGGATC
TCATGCTGGAGTTCTTCGCCCACGGGATCTCTGCGGAACAGGCGGTCGAAGGTGCCGATATC
ATTACGACAGCAACGGCCGACAAGCACAACGCCACGATCCTGAGCGACAATATGATCGGGCC
CGGCGTCCACATCAACGGCGTCGGCGGCGACTGCCCAGGCAAGACCGAGATGCACCGCGATA
TCTTGCTGCGTTCGGATATTTTCGTGGAGTTCCCGCCACAGACCCGGATGATCCCCGATCGT
TCAAACATTTGGCAATAAAGTTTCTTAAGATTGAATCCTGTTGCCGGTCTTGCGATGATTAT
CATATAATTTCTGTTGAATTACGTTAAGCATGTAATAATTAACATGTAATGCATGACGTTAT
TTATGAGATGGGTTTTTATGATTAGAGTCCCGCAATTATACATTTAATACGCGATAGAAAAC
AAAATATAGCGCGCAAACTAGGATAAATTATCGCGCGCGGTGTCATCTATGTTACTAGATCG
GGCCTCCTGTCAATGCTGGCGGCGGCTCTGGTGGTGGTTCTGGTGGCGGCTCTGAGGGTGGT
GGCTCTGAGGGTGGCGGTTCTGAGGGTGGCGGCTCTGAGGGAGGCGGTTCCGGTGGTGGCTC
TGGTTCCGGTGATTTTGATTATGAAAAGATGGCAAACGCTAATAAGGGGGCTATGACCGAAA

ATGCCGATGAAAACGCGCTACAGTCTGACGCTAAAGGCAAACTTGATTCTGTCGCTACTGAT
TACGGTGCTGCTATCGATGGTTTCATTGGTGACGTTTCCGGCCTTGCTAATGGTAATGGTGC
TACTGGTGATTTTGCTGGCTCTAATTCCCAAATGGCTCAAGTCGGTGACGGTGATAATTCAC
CTTTAATGAATAATTTCCGTCAATATTTACCTTCCCTCCCTCAATCGGTTGAATGTCGCCCT
TTTGTCTTTGGCCCAATACGCAAACCGCCTCTCCCCGCGCGTTGGCCGATTCATTAATGCAG
CTGGCACGACAGGTTTCCCGACTGGAAAGCGGGCAGTGAGCGCAACGCAATTAATGTGAGTT
AGCTCACTCATTAGGCACCCCAGGCTTTACACTTTATGCTTCCGGCTCGTATGTTGTGTGGA
ATTGTGAGaagCGGATAACAATTTCACACAGGAAACAGCTATGACCATGATTACGCCAAGCT
TGCATGCCTGCAGGTCCCCAGATTAGCCTTTTCAATTTCAGAAAGAATGCTAACCCACAGAT
GGTTAGAGAGGCTTACGCAGCAGGTCTCATCAAGACGATCTACCCGAGCAATAATCTCCAGG
AAATCAAATACCTTCCCAAGAAGGTTAAAGATGCAGTCAAAAGATTCAGGACTAACTGCATC
AAGAACACAGAGAAAGATATATTTCTCAAGATCAGAAGTACTATTCCAGTATGGACGATTCA
AGGCTTGCTTCACAAACCAAGGCAAGTAATAGAGATTGGAGTCTCTAAAAAGGTAGTTCCCA
CTGAATCAAAGGCCATGGAGTCAAAGATTCAAATAGAGGACCTAACAGAACTCGCCGTAAAG
ACTGGCGAACAGTTCATACAGAGTCTCTTACGACTCAATGACAAGAAGAAAATCTTCGTCAA
CATGGTGGAGCACGACACACTTGTCTACTCCAAAAATATCAAAGATACAGTCTCAGAAGACC
AAAGGGCAATTGAGACTTTTCAACAAAGGGTAATATCCGGAAACCTCCTCGGATTCCATTGC
CCAGCTATCTGTCACTTTATTGTGAAGATAGTGGAAAAGGAAGGTGGCTCCTACAAATGCCA
TCATTGCGATAAAGGAAAGGCCATCGTTGAAGATGCCTCTGCCGACAGTGGTCCCAAAGATG
GACCCCCACCCACGAGGAGCATCGTGGAAAAAGAAGACGTTCCAACCACGTCTTCAAAGCAA
GTGGATTGATGTGATATCTCCACTGACGTAAGGGATGACGCACAATCCCACTATCCTTCGCA
AGACCCTTCCTCTATATAAGGAAGTTCATTTCATTTGGAGAGAaagtACACGGGGGACTCTA
GAGGATCCCCGGGTACCGAGCTCGAATTTCCCCGATCGTTCAAACATTTGGCAATAAAGTTT
CTTAAGATTGAATCCTGTTGCCGGTCTTGCGATGATTATCATATAATTTCTGTTGAATTACG
TTAAGCATGTAATAATTAACATGTAATGCATGACGTTATTTATGAGATGGGTTTTTATGATT
AGAGTCCCGCAATTATACATTTAATACGCGATAGAAAACAAAATATAGCGCGCAAACTAGGA
TAAATTATCGCGCGCGGTGTCATCTATGTTACTAGATaaggCGGGAATTCACTGGCCGTCGT
TTTACAACGTCGTGACTGGGAAAACCCTGGCGTTACCCAACTTAATCGCCTTGCAGCACATC
CCCCTTTCGCCAGCTGGCGTAATAGCGAAGAGGCCCGCACCGATCGCCCTTCCCAACAGTTG
CGCAGCCTGAATGGCGCCCGCTCCTTTCGCTTTCTTCCCTTCCTTTCTCGCCACGTTCGCCG
GCTTTCCCCGTCAAGCTCTAAATCGGGGGCTCCCTTTAGGGTTCCGATTTAGTGCTTTACGG
CACCTCGACCCCAAAAAACTTGATTTGGGTGtATGGTTCACGTAGTGGGCCATCGCCCTGAT
AGACGGTTTTTCGCCCTTTGACGTTGGAGTCCACGTTCTTTAATAGTGGACTCTTGTTCCAA
ACTGGAACAACACTCAACCCTATCTCGGGCTATTCTTTTGATTTATAAGGGATTTTGCCGAT
TTCGGAACCACCATCAAACAGGATTTTCGCCTGCTGGGGCAAACCAGCGTGGACCGCTTGCT
GCAACTCTCTCAGGGCCAGGCGGTGAAGGGCAATCAGCTGTTGCCCGTCTCACTGGTGAAAA
GAAAAACCACCCCAGTACATTAAAAACGTCCGCAATGTGTTATTAAGTTGTCTAAGCGTCAA
TTTGTTTACACCACAATAtaattacatccctatattatttttctataaaagctgaaacttaa
attttatttattattgggattctatataactagctatattaatactcttttttgcttatactg
tataagcttgaaataaggacctattatttggtgcatttatttgactttattaatatcatcat
cactattactatattacgaattcctttttcttaagcttcaaaagcagaaaattaaataacat
agcacatagagatagtggttttattgctagcctaaattgcacaacacgaccacatcctaaat
cctaatgaagatggtcaaatttagactactttttttcttccattaagagtttataaatgtttg
atctgtatttgaatgcttttgttattgttactttagGTTAAAGTGTTTTTCGCTAGATTGGA
TGAAGAACTAAACAAAGTGAATCAGTTTCACAAGCCCAAAGAAACAGAGTTTTTGGAAAGAG
GAGAGATTCTGAAGAAACAGTTGGAGACTCTTGCAGAACTCAAACAGATCTTAAGTGATCGG
AAGAAGAGAAACTTGTCTGGCTCAAATTCACATCGCTCCTTCTCATCTTCTGTTCGAAACTC
TGATTTCTCTGCAGgttagtcttcttcttattaatgcttttactacaagttaggtcagattt
agggtactttagtaaattctgcattttgacttaagagttaagactcttattaaaagctaacc
attagtatatttgagatatgacgcggcctgcacccattagaaatcacaaaagtgcatgtcct
agaggccgtttacgtagtgcggctagtaaaaagaaaagtaggttatttaatatttttatatc
attgcacaatttagcgtttttgaaataatttaagtaaaaattcaaatgttttgttataacac

aatagaaattctactactgaatgacaagatttgttaagcatacaaatcttttgcatgtggat
gcaaaactcgaagctgcatatatgtgaaagaaagaattttagagttttgagattttattatg
tagacatgcatatcttcaatggccgacaagaacacaaatcgattctactaacaaacgaacaa
aatgtttgtgagtaagaaaataaaaatggaccgacaatagatggctaggtcgacaaaaattt
ttattttcttcttctctttagacctgtgttaatagtcacaaaatatataattatttgtaatt
tattttttcttattgttcagGGTCTCCAGGAGAACTAAGTGAGATACAAAGTGAAACATCAA
GAACAGATGAAATCATAGAGGCCTTAGAGAGGAACGGTGTGAGTTTCATAAACTCTGCAACG
AGGAGCAAAACAAAAGGAGGCAAACCAAAAATGTCTCTCCGCGTCGACATTCCCGACGCTGT
GGCCGGAGCCGAAGGTGGTATCGCAAGATCCATCGCCACCGCCATGTCTGTTCTTTGGGAAG
AGCTCGTTAACAACCCAAGATCAGATTTCACCAACTGGAAAAATATTCAAAGCGCCGAGAAG
AAGATACGAAGTGCCTTTGTTGAACTCTATAGAGGTCTTGGCTTGTTAAAGACTTACAGgta
tattcattaatgactcaatatcatttatttatttagtgatgatcttatcattttttaattctt
ttgttggctttattatggcagCTCGTTGAATATGATAGCTTTCACAAAAATAATGAAGAAAT
TCGATAAGgtaaattggttatagattgtactttcggtgataaaccaatgaaaaagataatca
tagtggttaatgatgaatctttgatatatatatgaatcatcagGTTGCTGGTCAGAATGCAT
CATCAACGTATCTCAAAGTCGTAAAGAGATCGCAATTTATCAGCTCTGATAAGgtaaaataa
aaggaggtcttatgctatgaataatttattgaaagattgtttgaaaatgattgtttgattaa
aattaaaaaagGTGGTAAGACTTATGGACGAAGTGGAGTCCATATTCACAAAGCACTTCGCC
AACAATGACAGGAAAAAGGCCATGAAATTCTTGAAACCCCACCAGACCAAAGATTCTCACAT
GGTCACTTTCTTTGTTGgtacttatttcatttctctatctttatacctttatataccattc
aaaaaaacagttcatcagagttttaacaaattgagattgtgtatctatgcagGGTTATTTAC
GGGTTGCTTCATCTCATTGTTTGTTATTTACATAATACTAGCCCATCTTTCTGGAATCTTCA
CTTCTAGTGATCAAGTCTCTTATCTGGAGACTGTTTATCCTGTTTTCAGgtaatgaataatt
atacgaattaatgatcaattcaacaaaactgtcaccatccaatgagacttaaccatttatcg
cttacattttgatgatttttttaaaaacgcagCGTTTTTGCGTTGCTGAGTCTACACATGT
TCATGTATGGATGCAATCTATACATGTGGAAGAACACGAGGATAAACTACACCTTTATTTTT
GAGTTTGCACCAAACACAGCGTTGCGTTACCGAGACGCGTTTCTGATGGGAACCACGTTCAT
GACCTCAGTTGTGGCAGCTATGGTCATCCACCTCATCCTCCGAGCCTCCGGTTTCTCAGCTA
GTCAAGTAGACACCATTCCAGGCATCCTCCTCCTGgtaaatcaaattacttagttcattaat
tatcatatggcgcgtttcaatcgcaatcgctatcacaatcacaatttgaaaccgctaatttc
tttttcggtgtgcatgctacagATCTTCATATGCGTCTTGATATGCCCATTTAACACATTCT
ACCGTCCAACAAGGTTCTGCTTCATCCGCATCTTGCGGAAGATTGTTTGCTCACCGTTCTAC
AAGgtaacaattggagttatttggttactttcagcacaagaaatagcagaaacatgattttt
ttttcttgtacggtaaatttagGTTTTGATGGTTGATTTCTTCATGGGCGATCAACTTACTA
GCCAGgtaaaactaagttatgcaacttcaatagatggtgacgacatctaatttagtcgataa
ttaacccattaatcgtgttctattcagATTCCATTGCTGAGACACCTTGAGACAACCGGGTG
TTACTTCTTGGCTCAAAGCTTCAAAACTCACGAATACAATACCTGCAAAAACGGAAGATACT
ATAGAGAATTTGCTTACTTGATTTCTTTCTTACCCTACTTCTGGCGTGCCATGCAAgtaagc
tcacttagggtttctctgtttttttttttttttgtcaagtcccttaaacctttcttctaa
gacactatgaacattaatttacagTGTGTAAGGAGATGGTGGGACGAATCAAACCCTGATCA
CCTAATCAACATGGGAAAATACGTGTCAGCGATGGTTGCAGCCGGAGTCCGCATAACCTACG
CGAGAGAAAACAACGACTTGTGGTTAACAATGGTGCTCGTAAGCTCCGTTGTGGCAACTATT
TACCAATTATACTGGGACTTTGTCAAGGATTGGGGTCTTCTAAACCCTAAATCGAAAAATCC
ATGGCTAAGAGACAATTTGGTTCTCCGGAACAAGAACTTCTACTACCTCTCCATTgtaagcc
aattacataactaactatagcgtgtttcacaatttatgatcttcgactaaatgttgagttgt
tcagGCGTTGAATTTGGTGTTGCGAGTTGCTTGGATCGAGACAATTATGAGATTCAGGGTCA
GTCCTGTTCAGTCTCATTTGCTAGATTTCTTCTTGGCGTCACTTGAAGTCATTCGTCGAGGC
CACTGGAACTTTTACAGgtaaataaaaaaacttcacctaggtttattaaaacttgattttgg
atgttattgaacatgaatctttctttcggtatttacagAGTGGAGAATGAGCACTTAAACAA
TGTCGGCCAATTTAGGGCAGTGAAGACCGTACCGTTACCGTTCCTTGACAGGGACTCAGACG
GTTAA

**PHO1 coding cDNA**

Sequence highlighted in green is deleted in *pho1-7* mutants (exon 2)

ATGGTGAAGTTCTCGAAGGAGCTAGAGGCACAACTTATACCGGAGTGGAAAGAGGCCTTTGTTAAC
TATTGTTTACTAAAGAAACAAATCAAGAAAATCAAAACCTCTCGTAAACCAAAACCGGCTTCTCATTA
CCCCATTGGTCATCACTCCGATTTTGGTCGATCTCTTTTCGACCCGGTTCGCAAATTGGCCAGGACCT
TCTCCGATAAACTATTTTCCAACTCAGAAAAACCAGAGATTCTCCAGGTAAGGAGAAGAAGAGGTA
GCTCAGAAACTGGGGATGACGTCGATGAGATTTACCAAACTGAACTTGTTCAGTTGTTTTCCGAAGA
AGACGAGGTTAAAGTGTTTTTCGCTAGATTGGATGAAGAACTAAACAAAGTGAATCAGTTTCACAA
GCCCAAAGAAACAGAGTTTTTGGAAAGAGGAGAGATTCTGAAGAAACAGTTGGAGACTCTTGCAG
AACTCAAACAGATCTTAAGTGATCGGAAGAAGAGAAACTTGTCTGGCTCAAATTCACATCGCTCCTT
CTCATCTTCTGTTCGAAACTCTGATTTCTCTGCAGGGTCTCCAGGAGAACTAAGTGAGATACAAAGT
GAAACATCAAGAACAGATGAAATCATAGAGGCCTTAGAGAGGAACGGTGTGAGTTTCATAAACTCT
GCAACGAGGAGCAAAACAAAAGGAGGCAAACCAAAAATGTCTCTCCGCGTCGACATTCCCGACGCT
GTGGCCGGAGCCGAAGGTGGTATCGCAAGATCCATCGCCACCGCCATGTCTGTTCTTTGGGAAGAG
CTCGTTAACAACCCAAGATCAGATTTCACCAACTGGAAAAATATTCAAAGCGCCGAGAAGAAGATAC
GAAGTGCCTTTGTTGAACTCTATAGAGGTCTTGGCTTGTTAAAGACTTACAGCTCGTTGAATATGATA
GCTTTCACAAAAATAATGAAGAAATTCGATAAGGTTGCTGGTCAGAATGCATCATCAACGTATCTCA
AAGTCGTAAAGAGATCGCAATTTATCAGCTCTGATAAGGTGGTAAGACTTATGGACGAAGTGGAGT
CCATATTCACAAAGCACTTCGCCAACAATGACAGGAAAAAGGCCATGAAATTCTTGAAACCCCACCA
GACCAAAGATTCTCACATGGTCACTTTCTTTGTTGGGTTATTTACGGGTTGCTTCATCTCATTGTTTGT
TATTTACATAATACTAGCCCATCTTTCTGGAATCTTCACTTCTAGTGATCAAGTCTCTTATCTGGAGAC
TGTTTATCCTGTTTTCAGCGTTTTTGCGTTGCTGAGTCTACACATGTTCATGTATGGATGCAATCTATA
CATGTGGAAGAACACGAGGATAAACTACACCTTTATTTTTGAGTTTGCACCAAACACAGCGTTGCGT
TACCGAGACGCGTTTCTGATGGGAACCACGTTCATGACCTCAGTTGTGGCAGCTATGGTCATCCACC
TCATCCTCCGAGCCTCCGGTTTCTCAGCTAGTCAAGTAGACACCATTCCAGGCATCCTCCTCCTGATC
TTCATATGCGTCTTGATATGCCCATTTAACACATTCTACCGTCCAACAAGGTTCTGCTTCATCCGCATC
TTGCGGAAGATTGTTTGCTCACCGTTCTACAAGGTTTTGATGGTTGATTTCTTCATGGGCGATCAACT
TACTAGCCAGATTCCATTGCTGAGACACCTTGAGACAACCGGGTGTTACTTCTTGGCTCAAAGCTTCA
AAACTCACGAATACAATACCTGCAAAAACGGAAGATACTATAGAGAATTTGCTTACTTGATTTCTTTC
TTACCCTACTTCTGGCGTGCCATGCAATGTGTAAGGAGATGGTGGGACGAATCAAACCCTGATCACC
TAATCAACATGGGAAAATACGTGTCAGCGATGGTTGCAGCCGGAGTCCGCATAACCTACGCGAGAG
AAAACAACGACTTGTGGTTAACAATGGTGCTCGTAAGCTCCGTTGTGGCAACTATTTACCAATTATAC
TGGGACTTTGTCAAGGATTGGGGTCTTCTAAACCCTAAATCGAAAAATCCATGGCTAAGAGACAATT
TGGTTCTCCGGAACAAGAACTTCTACTACCTCTCCATTGCGTTGAATTTGGTGTTGCGAGTTGCTTGG
ATCGAGACAATTATGAGATTCAGGGTCAGTCCTGTTCAGTCTCATTTGCTAGATTTCTTCTTGGCGTC
ACTTGAAGTCATTCGTCGAGGCCACTGGAACTTTTACAGAGTGGAGAATGAGCACTTAAACAATGTC
GGCCAATTTAGGGCAGTGAAGACCGTACCGTTACCGTTCCTTGACAGGGACTCAGACGGTTAA

**PHO1 protein sequence**

Sequence highlighted in red is deleted in *pho1-7* mutants
Sequence underlined form the 3 SPX subdomains, with amino acids in green involved in inositol polyphosphate binding

MVKFSKELEAQLIPEWKEAFVNYCLLKKQIKKIKTSRKPKPASHYPIGHHSDFGRSLFDPVRKLARTFSDKL
FSNSEKPEILQVRRRRGSSETGDDVDEIYQTELVQLFSEEDEVKVFFARLDEELNKVNQFHKPKETEFLERG
EILKKQLETLAELKQILSDRKKRNLSGSNSHRSFSSSVRNSDFSAGSPGELSEIQSETSRTDEIIEALERNGVS
FINSATRSKTKGGKPKMSLRVDIPDAVAGAEGGIARSIATAMSVLWEELVNNPRSDFTNWKNIQSAEKKI
RSAFVELYRGLGLLKTYSSLNMIAFTKIMKKFDKVAGQNASSTYLKVVKRSQFISSDKVVRLMDEVESIFTK
HFANNDRKKAMKFLKPHQTKDSHMVTFFVGLFTGCFISLFVIYIILAHLSGIFTSSDQVSYLETVYPVFSVFA
LLSLHMFMYGCNLYMWKNTRINYTFIFEFAPNTALRYRDAFLMGTTFMTSVVAAMVIHLILRASGFSAS
QVDTIPGILLLIFICVLICPFNTFYRPTRFCFIRILRKIVCSPFYKVLMVDFFMGDQLTSQIPLLRHLETTGCYFL
AQSFKTHEYNTCKNGRYYREFAYLISFLPYFWRAMQCVRRWWDESNPDHLINMGKYVSAMVAAGVRI
TYARENNDLWLTMVLVSSVVATIYQLYWDFVKDWGLLNPKSKNPWLRDNLVLRNKNFYYLSIALNLVLR
VAWIETIMRFRVSPVQSHLLDFFLASLEVIRRGHWNFYRVENEHLNNVGQFRAVKTVPLPFLDRDSDG
